# Distinct neural ensemble response statistics are associated with recognition and discrimination of natural sound textures

**DOI:** 10.1101/2020.01.30.914747

**Authors:** X. Zhai, F. Khatami, M. Sadeghi, F. He, H.L. Read, I.H. Stevenson, M.A. Escabí

## Abstract

The perception of sound textures, a class of natural sounds defined by statistical sound structure such as fire wind, and rain, has been proposed to arise through the integration of time-averaged summary statistics. Where and how the auditory system might encode these summary statistics to create internal representations of these stationary sounds, however, is unknown. Here, using natural textures and synthetic variants with reduced statistics, we show that summary statistics modulate the correlations between frequency organized neuron ensembles in the awake rabbit inferior colliculus. These neural ensemble correlation statistics capture high-order sound structure and allow for accurate neural decoding in a single trial recognition task with evidence accumulation times approaching 1 s. In contrast, the average activity across the neural ensemble (neural spectrum) provides a fast (tens of ms) and salient signal that contributes primarily to texture discrimination. Intriguingly, perceptual studies in human listeners reveals analogous trends: the sound spectrum is integrated quickly and serves as salient discrimination cue while high-order sound statistics are integrated slowly and contribute substantially more towards recognition. The findings suggest statistical sound cues such as the sound spectrum and correlation structure are represented by distinct response statistics in auditory midbrain ensembles, and that these neural response statistics may have dissociable roles and time scales for the recognition and discrimination of natural sounds.

**SIGNIFICANCE STATEMENT:** Being able to recognize and discriminate natural sounds, such as from a running stream, a crowd clapping, or ruffling leaves is a critical task of the normal functioning auditory system. Humans can easily perform such tasks, yet they can be particularly difficult for the hearing impaired and they challenge our most sophisticated computer algorithms. This difficulty is attributed to the complex physical structure of such natural sounds and the fact they are not unique: they vary randomly in a statistically defined manner from one excerpt to the other. Here we provide the first evidence, to our knowledge, that the central auditory system is able to encode and utilize statistical sound cues for natural sound recognition and discrimination behaviors.

## INTRODUCTION

What makes a sound natural, and what are the neural codes that support recognition and discrimination of real-world natural sounds? Although it is known that the early auditory system decomposes sounds into fundamental physical cues such as intensity, frequency and modulation, the higher-level neural computations that mediate natural sound recognition are poorly understood. This general lack of understanding is in part attributed to the structural complexity of natural sounds, which is difficult to study with traditional auditory test stimuli, such as tones, noise or modulated sequences. Such stimuli can reveal details of the neural representation for relatively low-level acoustic cues, yet they don’t capture the rich and diverse statistical structure of natural sounds. Thus, they cannot reveal the computations associated with higher-level sound properties that facilitate auditory tasks such as natural sound recognition or discrimination. A class of stationary natural sounds termed *textures*, such as the random sounds emanating from a running stream, a crowded restaurant, or a chorus of birds, have been proposed as alternative natural stimuli which allow for manipulating high-level acoustic structure ^1^. Texture sounds are composed of spatially and temporally distributed acoustic elements that are collectively perceived as a single source and are defined by their statistical features. Identification of these natural sound has been proposed to be mediated through the integration of time-averaged *summary statistics*, which account for high-level structures such as the sparsity and time-frequency correlation structures of natural sounds ^1-3^. Using summary statistics from a generative model of the peripheral and central auditory system, it is possible to synthesize highly realistic auditory texture sounds ^1^. This suggests that high-order statistical cues are perceptually salient and that the brain might extract these statistical features to build internal representations of sounds.

Where and how the brain represents summary statistics in neural activity and how these contributes to recognition and discrimination for real world sounds is largely unknown. Neural activity throughout the auditory pathway is known to be modulated by a variety of statistical cues such as the sound contrast, modulation power spectrum, and correlation structure ^4-12^. However, it has been less clear whether and to what degree summary statistics serve as informative stimulus dimensions that directly contribute to sound recognition. The inferior colliculus (IC) is one candidate mid-level structure for representing such summary statistics. As the principal midbrain auditory nucleus, the IC receives highly convergent brainstem inputs with varied sound selectivities. Neurons in the IC are selective over most of the perceptually relevant range of sound modulations and neural activity is strongly driven by multiple high-order sound statistics ^4-7,10^. In previous work, we showed the correlation statistics of natural sounds are highly informative about stimulus identity and they appear to be represented in the correlation statistics of auditory midbrain neuron ensembles ^4^. Correlations between neurons have also been proposed as mechanisms for pitch identification ^13^ and sound localization ^14^. This broadly support the hypotheses that high-order sound statistics are reflected in the response correlations of neural ensembles and that these correlations could potentially sub-serve basic auditory tasks.

Here using natural and synthetic texture sounds, we test the hypothesis that statistical structure in natural texture sounds modulate the response statistics of neural ensembles in the IC of unanesthatized rabbits, and that neural response statistics can directly mediate sound recognition and discrimination behavior. By comparing the performance of neural decoders with human texture perception we find that low-level spectrum-based cues are integrated on relatively fast time scales (tens of ms) and appear to contribute primarily towards texture discrimination. High-order statistical cues, by comparison, require substantially longer evidence accumulation times (> 500 ms) and contribute substantially more towards sound recognition. Collectively, the findings suggest that spectrum cues and accompanying place rate representation (neural spectrum) contribute surprisingly little towards the recognition of auditory textures. Instead, high-order statistical sound structure is reflected in the distributed patterns of correlated activity across IC neural ensembles and such neural response structure contributes substantially more towards the recognition of natural auditory textures.

## RESULTS

### Natural Sound Texture Statistics Modulate Neural Correlation but not Neural Spectrum Statistics

To determine how natural sound statistics influence the response statistics of neural ensembles in IC, we first characterized several key statistics from an auditory model representation for five natural sound recordings. These include sounds from a crackling fire, bird chorus, outdoor crowd, running water, and a rattling snake (Supplementary sounds 1-5). We then used texture synthesis ^1^ to generate synthetic sound variants with perturbed low- and high-order statistics (see METHODS). The selected sound textures have distinct spectral and temporal properties, and they each show diverse structures in the measured statistics (Fig. 1). The synthetic sound variants are generated by sequentially imposing the cochlear channel power statistics (i.e., power spectrum; Fig. 1A and B; Supplementary sounds 6-10), cochlear channel marginal statistics (envelope mean, variance, and skew; Fig. 1B; Supplementary sounds 11-15), the modulation power of each cochlear channel (Fig. 1C; Supplementary sounds 16-20), and the correlation structure between cochlear (Fig. 1D; Supplementary sounds 21-25) and modulation channels. As illustrated, there are marked differences in both low- and high-order statistical cues across the different natural sounds. For instance, the crowd and water sounds have relatively similar power spectra (Fig. 1A and B) and modulation power (Fig. 1C), and the correlations between frequency channels are weak, as reflected in the diagonalized correlation matrix (Fig. 1D). The rattling snake sound, by comparison, has a power spectrum that is biased towards higher frequencies (Fig. 1A and B). Furthermore, the cochlear channels are highly correlated across frequencies (Fig. 1D) with modulation power concentrated at about 20 Hz (Fig. 1C), which reflects the coherent periodicity of the broadband rattling sound. Collectively, the differences in statistical structure for these natural textures could differentially drive neural responses in IC and may contribute towards recognition or discrimination of these sounds.

**Fig. 1:**
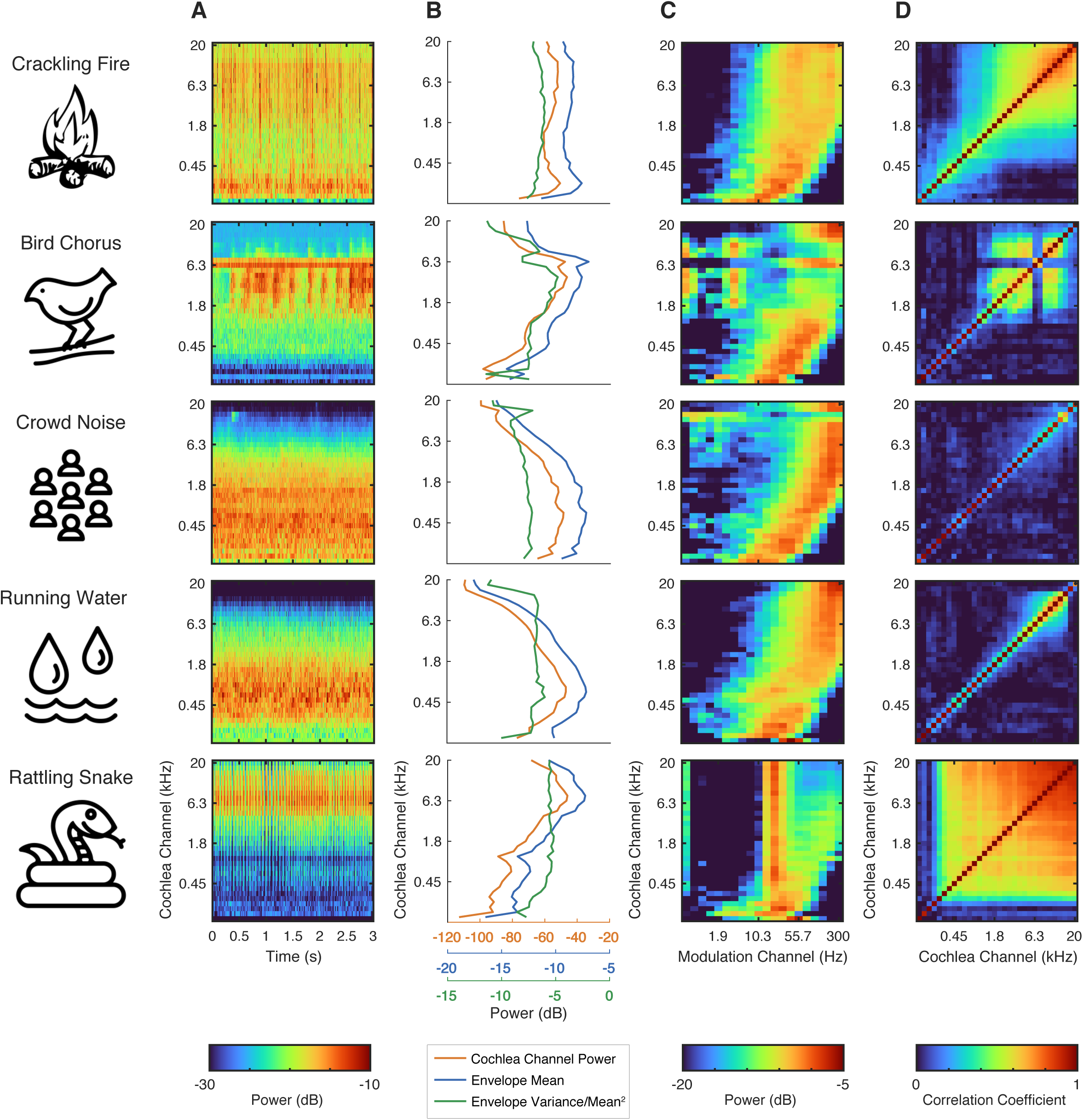
Sound summary statistics measured from five natural sound textures. (A) Cochlear model spectrograms. Color scale indicates the time-varying envelopes decomposed from frequency-organized cochlear filters. (B) Cochlear channel power and envelope marginal moments (mean and variance/mean^2^). (C) Modulation power spectrum. The power of each modulation band is normalized by the variance of the corresponding cochlear envelope and plotted as a function of modulation frequency (Hz). (D) Cochlear cross-band envelope correlations.

Though human listeners are perceptually sensitive to statistical cues in natural sound textures, there is little evidence on how neural responses to these statistics may contribute to texture perception. Here, we used the synthetic and original texture sounds as stimuli to determine whether sound statistics modulate the response statistics of neural ensembles in the unanesthetized rabbit IC (*N*=4 animals, 29 penetration sites were included for analysis). Fig. 2 demonstrates the neural response statistics measured from a representative penetration site for a synthetic bird sound (synthesized variant that includes all the statistics) using an analog representation of multi-unit activity (aMUA, see METHODS). From the response neurogram (Fig. 2B, average activity across response trials) we estimated the stimulus-driven **neural correlation** statistics by correlating the signals from all electrode channel pairs across independent response trials (see METHODS). Fig. 2F and G illustrate different degrees of correlated neural activity for the aMUA signals from example recording channel pairs. The neural responses from channel 7 and 8, for instance show a significantly correlated temporal signature (Fig. 2F; p<0.01, Fisher z-transform test) which is reflected in the point wise scatter plot and ultimately in the measured correlation coefficient (r=0.89) and stimulus driven correlation (c=0.286). Similarly, channel 7 shows a significant although weaker correlation with channels 11 (r=0.33, c=0.11) and 16 (r=0.16, c=0.05). Since our recordings from multiple spatially separated electrode channels follow the tonotopic ordering of the IC (Fig. 2A, frequency response areas cover a frequency range of approximate 0.5-10 kHz for this penetration site), we refer to the cross-channel correlations at zero lag as **spectral correlations** (Fig. 2C). Conceptually, this metric captures the degree to which distinct neural recording channels are temporally synchronous with one another ^4,15^, analogous to the model-based channel correlation ^16^ (Fig. 1D). We also estimated the neural response correlations across time for each neural recording channel which we refer to as the **temporal correlations** (Fig. 2D; see METHODS). As previously shown, the temporal correlations capture the stimulus-driven temporal response pattern for each individual channel^4^ and is closely related to the modulation spectrum of the sound (related via a Fourier Transform) ^1^. Both the spectral and temporal correlations shown here are “stimulus” driven correlations, where “noise” correlations on single trials have been removed by trial shuffling (see METHODS). Finally, for each recording location, we also measured the **neural spectrum** statistic assessed by computing the average response amplitude from each electrode channel over time and across response trials (Fig. 2E, see METHODS). This neural response statistic closely resembles the model-based sound spectrum which is widely used to measure the frequency composition of sounds.

**Fig. 2:**
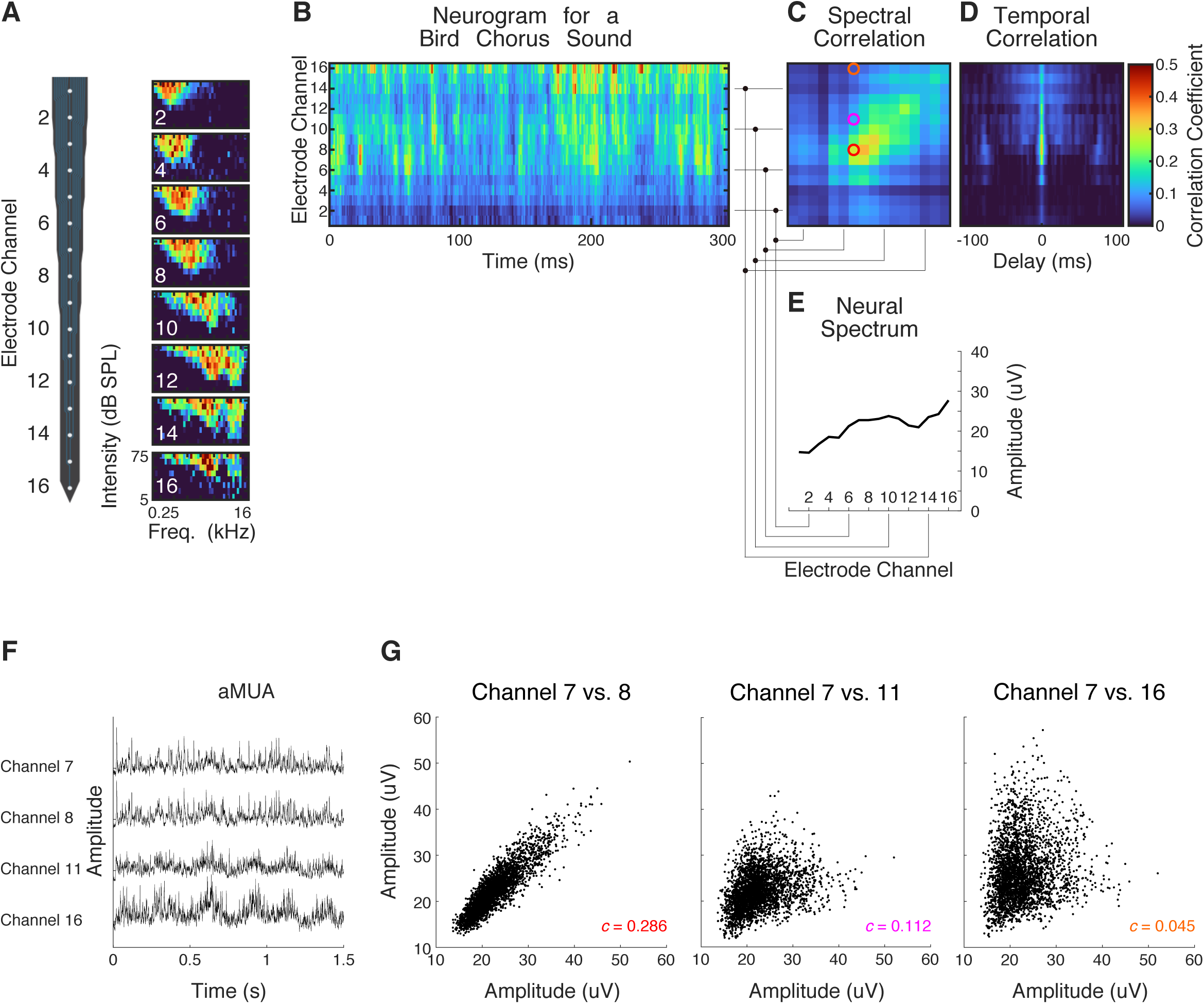
Measuring neural response statistics from auditory midbrain ensembles. (A) A linear electrode array used for neurophysiological recordings. Frequency response areas are shown for eight of the recording channels in a representative tonotopically organized inferior colliculus (IC) penetration site (red indicates high activity and blue indicates low activity). (B) A segment of analog multiunit activity (aMUA) neural signals across 16 recording channels (red indicates strong response and blue indicates weak response) for a synthetic bird chorus sound texture (full statistics condition, see METHODS). (C-D) Stimulus-driven spectral and temporal neural ensemble correlations obtained from the aMUA activity. Spectral correlations are obtained by correlating the aMUA signals for different channels while temporal correlations are obtained by computing autocorrelations of individual channels (see METHODS). (E) The neural spectrum is calculated from the aMUA by computing the mean response for each of the 16 channels. (F) aMUA signals for 4 example recording channels from (B). (G) Scatter plots using the aMUA signals in (F) for three different channel pairs illustrate different degrees of correlation. The corresponding pixels in the spectral correlation matrix shown in (C) have the stimulus-driven correlation coefficients of 0.286, 0.112, and 0.045, respectively.

If higher order statistics of sound provide meaningful cues for identifying sounds, neural responses should reflect and vary systematically with statistical variation of natural sounds. As seen in the example of Fig. 1, high-order natural sound statistics vary markedly for each natural texture, which could provide useful information about the identity of the sound. The measured neural correlations and neural spectrum varied markedly across the five texture sounds and across synthetic variants with different statistics as seen for the penetration site of Fig. 2 (Fig. 3; additional examples in Supplementary Fig. 1). In general, spectral (Fig. 3A) and temporal (Fig. 3B) correlations to different natural sound textures are highly diverse and reflect stimulus-dependent structures. For example, in the synthetic fire sound containing the full set of statistics (Fig. 3 column of +Corr. condition), spectral correlations are extensive with the envelopes of both nearby and distant electrode channels showing correlated activity. By comparison, the neural correlations to the crowd sound are localized to neighboring channels with relatively low frequencies, and the response to the snake sound exhibits strong correlated activity between channels with best frequencies above approximately 1 kHz. Temporal correlations also show distinct, stimulus-dependent patterns. The temporal correlations of the bird sound, for instance, show a broad/slow component at high frequencies, while the correlations of fire, crowd, and water sounds are narrow/fast and show little frequency selectivity. In contrast to all four other sounds, the snake sound exhibits periodic correlations for high frequency channels (∼50 ms period) that reflect the periodic structure of snake rattling at approximately 20 Hz. The neural spectra (Fig. 3C) shows a somewhat lower amount of diversity across sounds. The fire, bird and snake sounds induce the strongest activity for high frequency channels, while crowd and water more strongly drive low frequency channels. Thus, neural response correlations reflect statistical structure that can potentially distinguish each of the natural texture sounds.

**Fig. 3:**
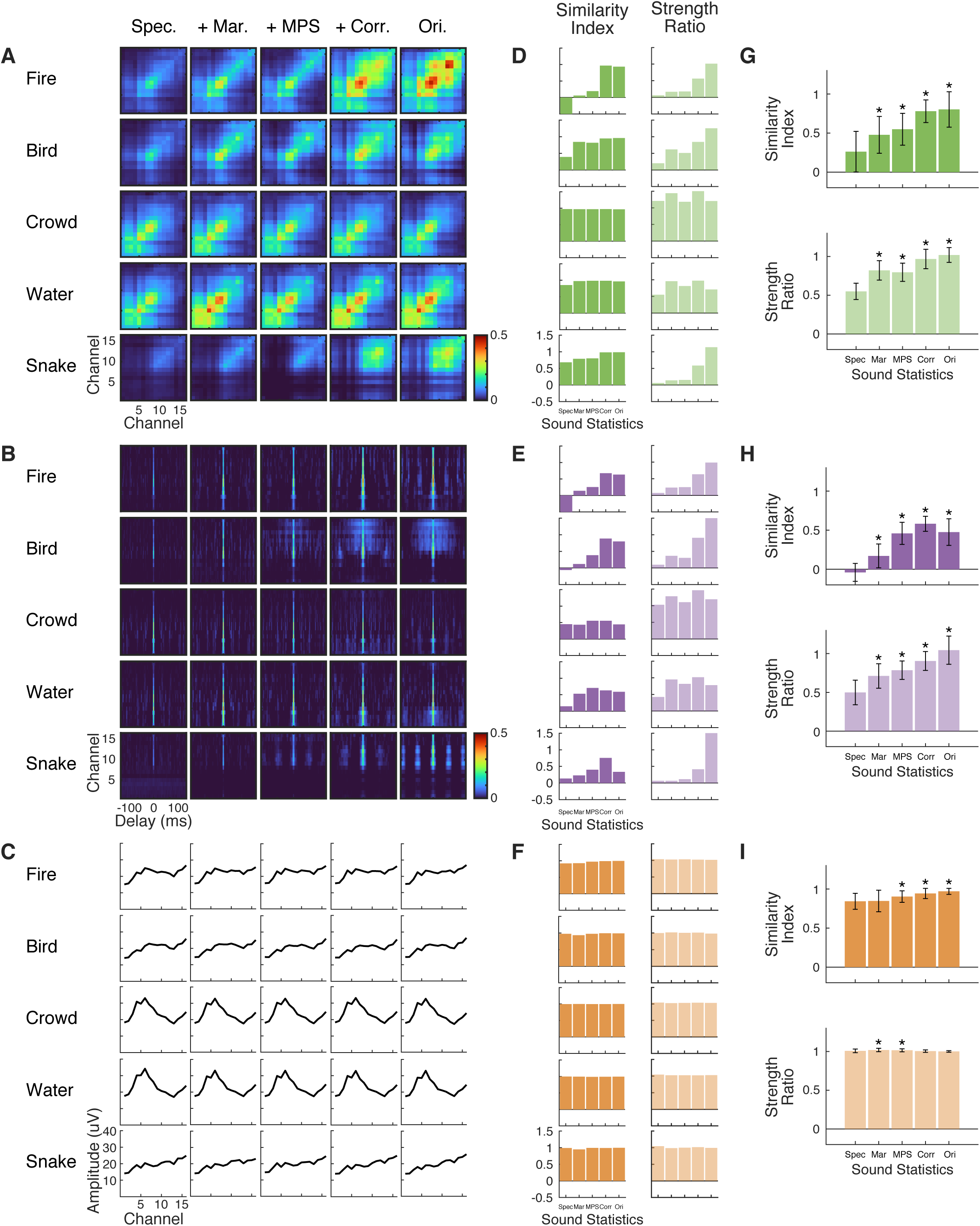
Neural correlations and neural spectrums for synthetic variants and original sounds. (A) Spectral correlations for the IC penetration site shown in Fig. 2 (same site for B-F). (B) Temporal correlations for the same recording site. Spectral and temporal correlations are shown for the original sound (Ori) and synthetic variants with different statistics included (Spec., +Mar., +MPS, +Corr.). Neural correlation matrices show distinct structures and unique patterns across the five tested sounds, while for each sound, such stimulus-dependent correlations converge upon adding sound statistics. (C) Neural spectrums. Although the neural spectrums show diversity across sounds, they do not change significantly with sound statistics included. (D-F) Similarity index and strength ratio as a function of sound statistics for three response metrics. Comparisons are made between the response of a synthetic sound variant and that of the original sound. For spectral (D) and temporal (E) correlations, both similarity index and relative strength increase upon adding sound statistics to synthetic variants of fire, bird and snake sounds. For neural spectrums (F), the similarity index and relative strength are relatively constant with different statistics for all sounds. (G-H) Average similarity index and strength ratio as a function of sound statistics across *N*=29 penetration sites (*N*=4, 9, 8, 8 from four animals). Averaged across five sounds, the similarity index and strength ratio of spectral (G) and temporal (H) correlations increase with statistics, indicating the correlation matrices converge to the original sound structure upon adding sound statistics. The neural spectrum (I) similarity index and strength ratio do not show a significant change with added sound statistics. Error bars, S.D. Asterisks indicate a statistical significance increase when compared to the Spec. condition (*p* < 0.05, paired t-test, corrected for multiple comparisons). Spec.: cochlear spectrum; Mar.: cochlear marginals; MPS: modulation power spectrum; Corr.: correlation; Ori.: original in this and all subsequent figures.

In addition to the differences in responses statistics observed for each texture sound, neural correlation statistics in the IC also varied systematically with the sound statistics that were included in the perturbed texture variants. Adding sound statistics to the synthetic variants increases the strength of spectral and temporal correlations between neural responses. Moreover, the patterns of neural correlations change and become increasingly similar to the original sound responses as statistics are added (Fig. 3A and B). The same was not true for the neural spectrum, which was largely unaffected by adding high-order statistics to the synthetic sound variants and consistently resembled the response to the original sound (Fig. 3C). To quantify this effect, we calculated two indices that compare the neural correlations and spectrum for the reduced and original sound variants: a cross-validated similarity index (SI) and strength ratio (SR) (see METHODS). In the example penetration site, the SI and SR of the spectral (Fig. 3D) and temporal (Fig. 3E) correlations for fire, bird and snake sounds increase as more statistics are included in the sound variants. Thus, the correlation strength increases upon adding high-order statistics while the correlation pattern converges on that of the original sound. By comparison, for crowd and water sounds, the neural correlations do not change substantially with the included sound statistics, indicating that the neural correlation statistics are similar to the response of original sounds even when only the sound spectrum is imposed. Such lack of sequential change for these sounds reflects the fact that the crowd and water sounds are both well characterized by relatively low-level acoustic structure. They have minimal acoustic correlations and are relatively random over time (Fig. 1). By comparison, although the neural spectrum shows a fair amount of diversity across sound textures, it is very stable regardless of which statistics are included in the synthetic variants (Fig. 3F). Consequently, the measured SI and SR of the neural spectrum for this recording site are near-constant for all five sounds.

Similar trends are observed across the neural population indicating that high-order statistics of the sounds strongly influence the neural ensemble activity (Fig. 3G-I; for individual sounds, see Supplementary Fig. 2). Neural correlations change systematically and converge on that of the original sound upon adding sound statistics, whereas neural spectrum is relatively stable and resembles the original sound condition regardless of which statistics are included in the reduced sounds (Fig. 3G-I). Averaged across all five sounds and penetration sites, the SI of spectral correlations shows a significant increase from 0.26 ± 0.26 (mean ± SD), when only the sound spectrum is included (i.e., Spec. condition), to 0.78 ± 0.14, when the marginals, modulation power and correlations are included (i.e., + Corr condition) in the synthetic sound stimuli (*p* < 0.05, paired t-test comparing each condition against Spec; corrected for multiple comparisons, applied to this and all subsequent t-tests). Similarly, the SI of temporal correlations increases systematically from −0.04 ± 0.11 to 0.58 ± 0.09 (*p* < 0.05, paired t-test). The SR of the neural correlations statistics also systematically increased upon adding statistics to the synthetic variants (spectral = 0.55 ± 0.10 to 0.97 ± 0.13; temporal = 0.50 ± 0.16 to 0.90 ± 0.12; *p* < 0.05, paired t-test). These results differ from those for the neural spectrum, which showed only a slight increase in SI (from 0.84 ± 0.10 for Spec. to 0.94 ± 0.06 for +Corr.; *p*<0.05, paired t-test) whereas the SR remains near-constant as statistics are added (∼1, all S.D. < 0.02; N.S., paired t-test). Overall, these findings demonstrate that neural correlations in IC become stronger and their pattern converges on that of the original sound textures upon adding high-order statistics while the neural spectrum shows substantial less variation and is largely unaffected by high-order sound statistics.

### The Contribution of Neural Response Correlation and Spectrum Statistics to Recognition and Discrimination of Texture Sounds

Given that the neural response pattern and strength is strongly modulated by the high-order texture statistics, we next explored how the response spectrum- and correlation-based neural codes can contribute to recognition and discrimination of sound textures. We used a single-trial neural decoder (see METHODS), to determine whether the neural correlations and spectrum in IC could allow sound textures to be identified and discriminated. In the **recognition task**, a naïve Bayes classifier was trained with the neural spectrums or correlations of the five original sounds (obtained with 1000 ms window). The classifier was required to identify the delivered sound using single response trials of a variable duration (62.5 to 1000 ms in half-octave steps). Fig. 4A shows the cross-validated (half of the data was used for training the model and the other half for validation, see METHODS) classification performance for the example penetration site shown previously (Fig. 3) when the stimuli were synthetic sounds that included all statistics (+Corr. condition, applied to Fig. 4A-D; results for all other statistic conditions are shown in Supplementary Fig. 3). At short durations, the performance of neural correlation-based classifiers ranges from approximately 0% to 90% for different sounds, although the average performance is above chance (32.2%, 29.2%, and 32.2% for spectro-temporal, spectral, and temporal, respectively at 62.5 ms; chance is 20%). For most sounds, classifier performance increases with sound duration. An exception to this, was the temporal classifier performance for the snake sound, which was below chance. This misclassification was likely due to the fact that the rattling frequency between the first and second half of the data was different (approximately 15 vs. 25 Hz) which affected the cross-validation results (see METHODS). In contrast to the correlation-based classifiers, where performance tends to improve with sound duration, the neural spectrum classifier shows relative stable and high performance even for short sound durations (67.0% at 62.5 ms).

**Fig. 4:**
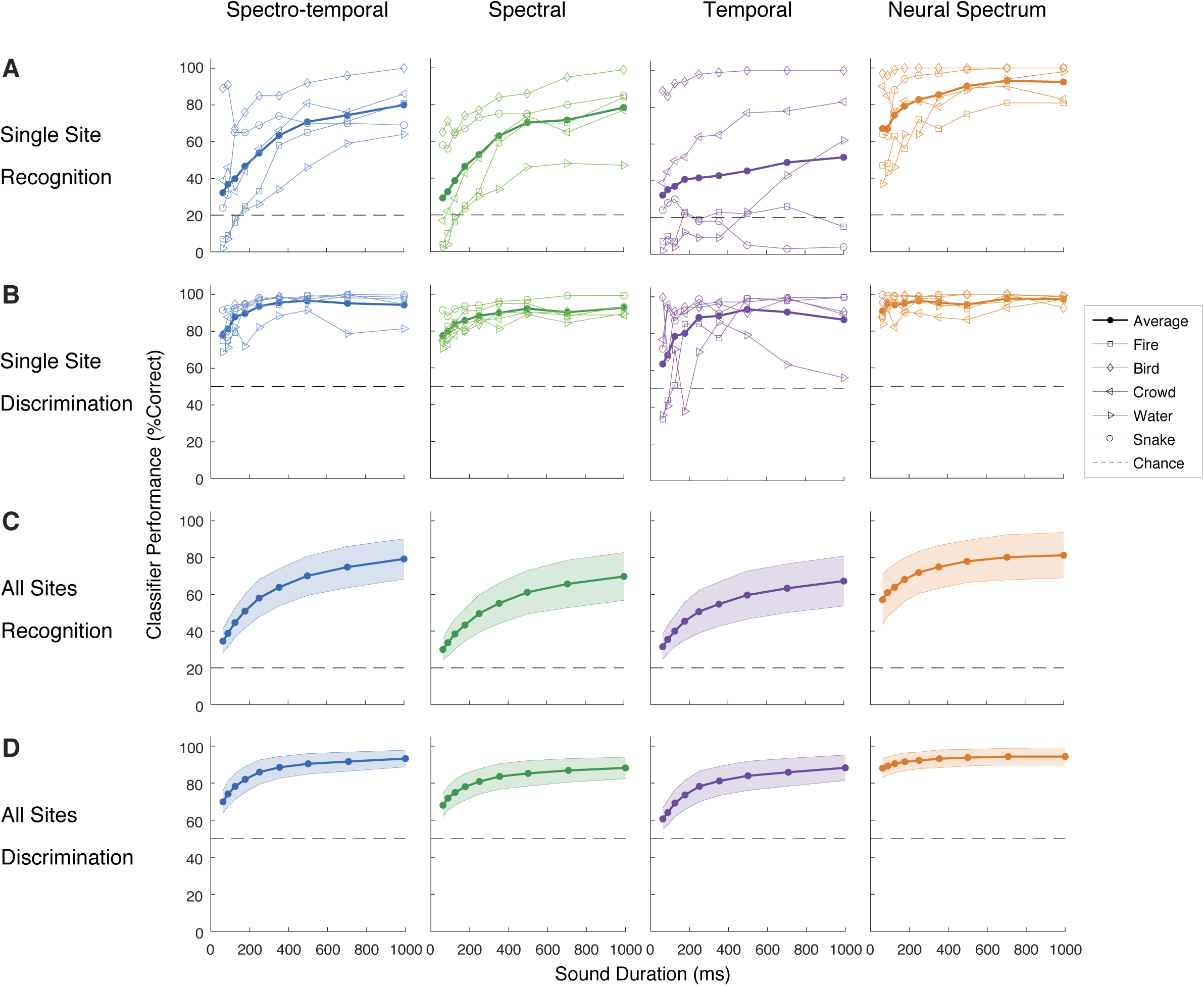
Decoding neural correlations and neural spectrums for sound recognition and categorization. Classification results are shown as a function of sound duration. Stimuli were synthetic sounds with full statistics (+Corr. condition, see METHODS). (A) Single-trial classification results for the recognition task (see METHODS) for the penetration site shown in Fig. 3. The performance of each individual sound (black curves) and the average results (red curve) are shown for different classifiers (spectro-temporal, spectral, temporal, and neural spectrum). (B) Classification results for the discrimination task for the same penetration site in (A). (C-D) Average performance across *N*=29 penetration sites. Shaded areas, S.D.

Similar results are observed across all penetration sites (Fig. 4C). The performance of all classifiers is above chance and increases with sound duration. Among the correlation-based classifiers, the spectro-temporal classifier shows the highest performance (increase from 34.6% to 79.3% with duration), followed by the spectral (30.0% to 69.7%) and the temporal (31.5% to 67.3%) classifiers. In contrast, the neural spectrum classifier has more stable and higher performance (57.1% to 81.3%) that does not improve substantially with sound duration (a ∼25% increase compared to 40%-45% for the correlation-based classifiers). Thus, spectrum- and correlation-based neural codes can both contribute to texture recognition, and performance tends to improve with the sound duration.

Although texture recognition performance of the synthetic texture (+Corr. condition) depends strongly on the sound duration and neural response statistics measured, texture discrimination was less dependent on both. In the texture **discrimination** task, the naïve Bayes classifier was trained using the responses for sound pairs of identical duration (see METHODS). For the example penetration site of Fig. 4B, most sounds can be easily discriminated regardless of the sound duration used. One exception is for the water sound where, for this example, the temporal correlation-based classifier performance tends to decrease with duration, although it remains above chance (50%). Averaged across all sounds and all penetration sites (Fig. 4D), performance is high and above chance for all classifiers when the sound is 62.5 ms (spectro-temporal 70.0%, spectral 68.1%, temporal 60.1%, neural spectrum 88.0%) and shows slight increases with sound duration. The performance reaches and exceeds 90% with longer durations.

To further evaluate whether and the degree to which the sound spectrum itself may be driving correlated neural activity that might contribute towards the recognition and discrimination of textures, we repeated the experiments in *N*=11 recording sites (from 2 animals) using the original sound textures and texture variants with an equalized 1/f power spectrum ^4^ (Supplementary sounds 26-30). This manipulation guarantees that average spectrum cues are the same across sounds while preserving many of the high-order sound cues in the original sounds. Despite the fact that these sounds have identical spectrum, the sounds are perceptually distinct and are uniquely perceived as the original sound. Here, the neural correlations of the equalized sounds are distinct from each other and remarkably similar to the original sounds (Fig. 5A, spectral; Fig. 5B, temporal). On the other hand, while the neural spectra of the original sounds were quite distinct, the neural spectra for the equalized sounds are all quite similar to each other (Fig. 5C). Thus, although the neural spectrum is strongly affected by the sound spectrum, high-order structure in these equalized sounds appears to be largely preserved and encoded by the neural correlations within IC ensembles.

**Fig. 5:**
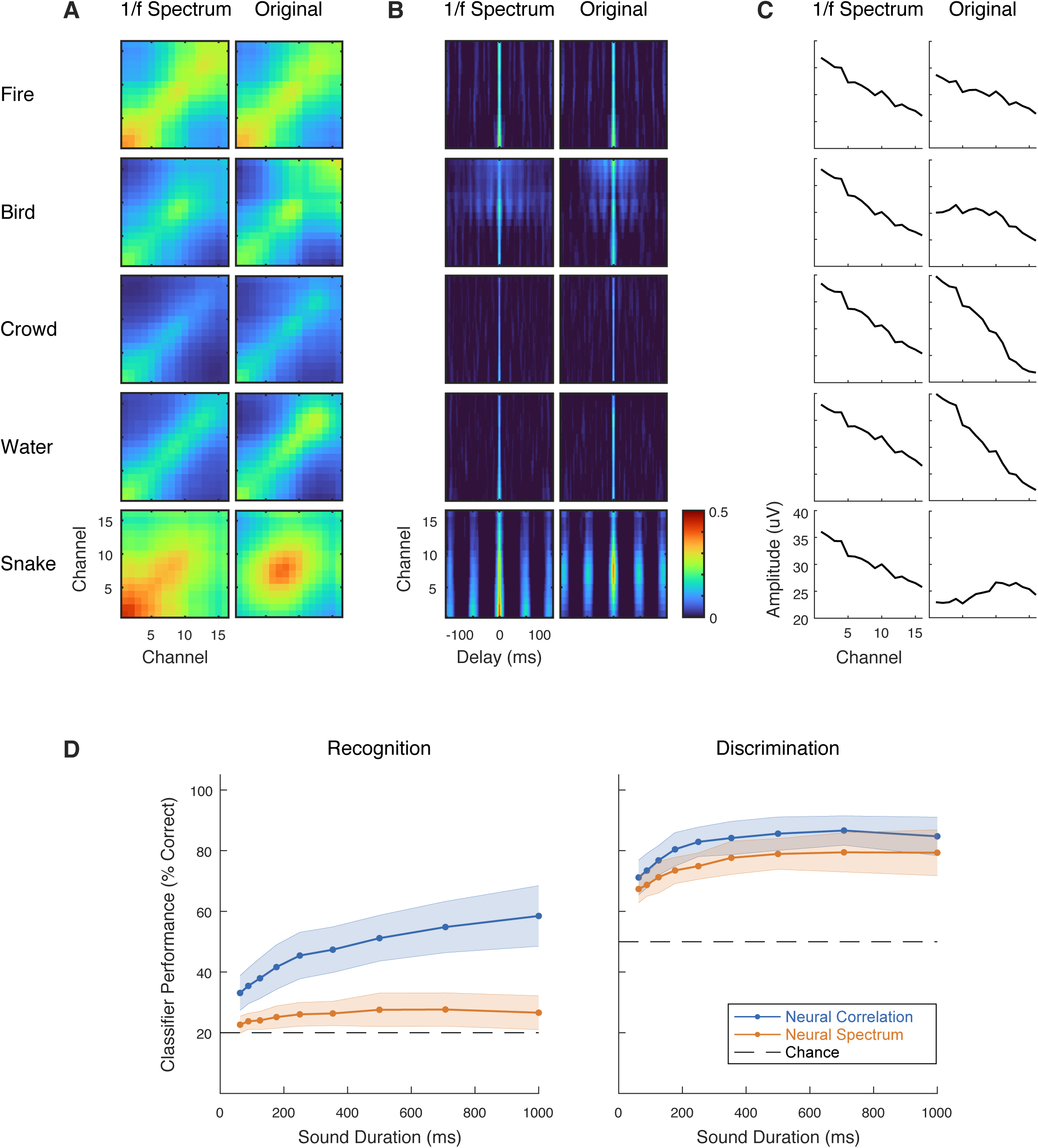
Neural correlations and neural spectrums for sound variants with equalized power spectrum. (A-C) Response metrics of a single IC penetration site. Spectral (A) and temporal (B) neural correlations show similar structures between the 1/f equalized and original sounds. Neural spectrums (C) show nearly identical structures across sounds for the 1/f condition while they are different from those in the original sounds. (D) Average single-trial classification performance across *N*=11 penetration sites (*N*=3, 8 from two animals). In the recognition task, the neural classifier is trained using the responses to the original sounds, while the validation data are from the spectrum-equalized sounds. In the discrimination task, both training and testing data are from the responses to the spectrum-equalized sounds (see METHODS).

How does the removal of the sound spectrum affect classification performance? Fig. 5D shows the population average neural identification and discrimination for the spectrum equalized sounds. While the neural spectrum contributed substantially to recognition of both the synthetic and original sounds (Fig. 4C, far right; Supplementary Fig. 3A, Ori. far right) neural spectrum identification performance drops to near chance (from 90.1% to 26.6% at 1 s) for the spectrum equalized sounds (Fig. 5D, red). Recognition performance for the neural correlation also drops compared to the original sounds (from 84.9% to 58.5% at 1 s) but still retains information for the spectrum equalized sounds that allows the classifier to perform well above chance (Fig. 5D, blue). Similar results are also seen for the discrimination task. The neural correlation classifier performance is only slightly reduced compared to the original sound (93.6% vs. 84.7% at 1 s), however, the neural spectrum classifier has a larger reduction in performance (96.0% to 79.3%). These findings suggest that high-order sound structure from the original sounds is retained in the neural correlations despite equalization and that this structure can contribute to neural recognition and discrimination independently of the cues in the sound spectrum.

Next, we explored how the addition of high-order sound statistics affects how well sound textures can be identified or discriminated from neural responses. The neural classifier performance at 1 s duration is shown in Fig. 6A as a function of the sound statistics that were included during the synthesis. For the recognition task, the performance of all classifiers increases substantially upon adding sound statistics (for spectral classifier and temporal classifier, see Supplementary Fig. 4). The spectro-temporal classifier shows the highest performance amongst the correlation-based classifiers, increasing from 40.0 ± 11.6% for the Spec. condition to 83.0 ± 9.4% for Ori. This was followed by the spectral correlation-based classifier (39.2 ± 11.8% to 78.1 ± 10.2%) and the temporal correlation-based classifier (25.2 ± 6.0% to 63.6 ± 11.8%), which showed the lowest performance on average. In contrast to the correlation classifiers, the neural spectrum classifier exhibited very high performance that was much less dependent on the included statistics. The recognition accuracy is 66.6 ± 14.7% for the Spec. condition and improves to 84.3 ± 10.7% for Ori condition. Note that the spectro-temporal and neural spectrum classifiers have roughly similar high performance (about 80%) for the synthetic sounds with correlation statistics and original sounds. However, in the discrimination task, the performance doesn’t improve substantially with sound statistics for the spectro-temporal, spectral, and neural spectrum classifiers, and improves only slightly (∼18%) for the temporal classifier. Together, these findings suggest that correlation- and spectrum-based cues can contribute differently to neural recognition and discrimination of texture sounds.

**Fig. 6:**
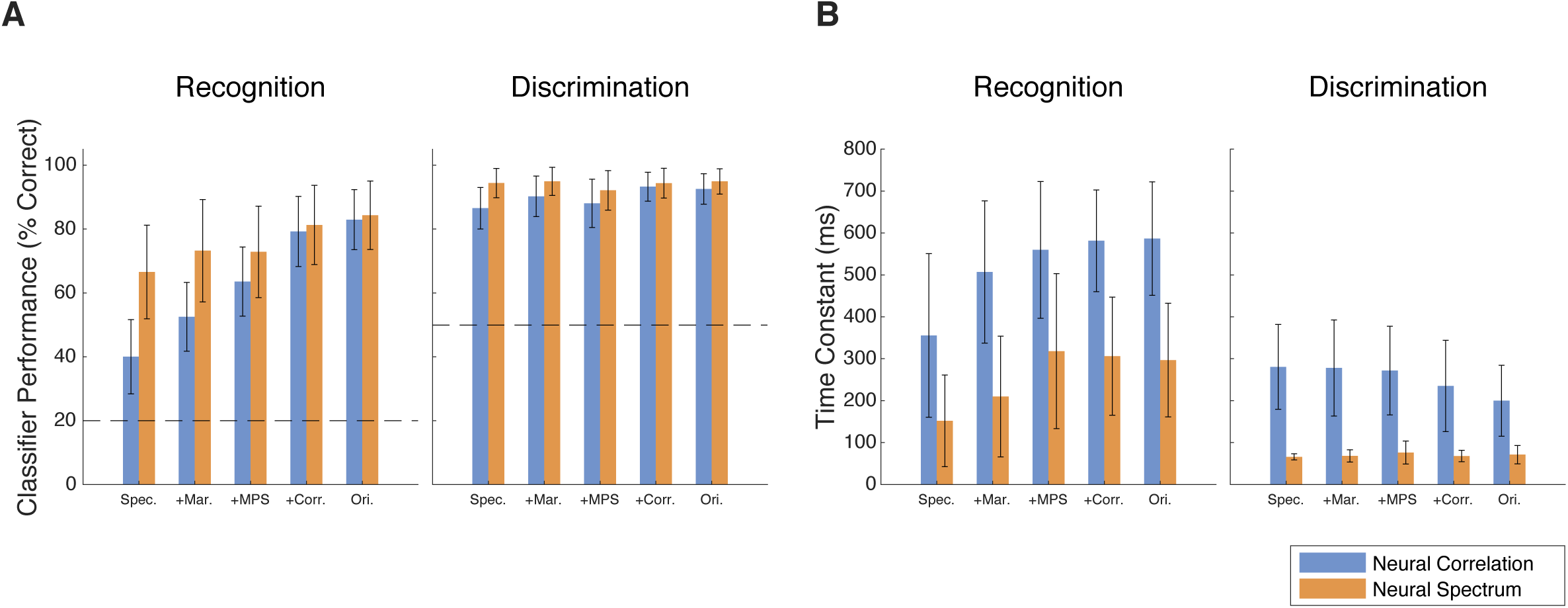
Neural classifier performance and time constant as a function of sound statistics. (A) Classification results for the spectro-temporal and neural spectrum classifiers in the recognition and discrimination tasks. The average performance is shown for 1 s sound duration (averaged across sounds and penetration sites). Recognition performance improves substantially upon adding sound statistics to the synthetic variants. Perofrmance for the discrimnation task is much more stable and does not improve substantially with added statistics. (B) Time constant (time required to reach 90% of the maximum measured performance) as a function of sound statistics in the recognition and discrimination tasks. Time constants for the recognition task are substantially longer than the discrimnation task. Error bars, S.D.

The evidence accumulation times for the neural correlation and spectrum decoders differ significantly as a function of task (recognition vs discrimination), the statistics that are included in the synthetic texture stimuli, and the neural response statistics used for classification (neural correlation vs. spectrum) (p<0.001, m-way ANOVA). Fig. 6B shows the time constant, defined as the time it takes to reach 90% of the corresponding maximum classifier performance, as a function of sound statistics. First, for a both the spectrum and correlation classifiers, time constants for the discrimination task are shorter than the recognition task and they exhibit different behaviors as a function of added sound statistics. In the recognition task, the time constant increases systematically with additional sound statistics (change between conditions Spec. to Ori: 65.4% increase, from 355 ± 195 ms to 587 ± 135 ms for correlation; 95.4% increase, from 152 ± 109 ms to 297 ± 135 ms for neural spectrum). This systematic increase in the time constant is partly attributed to performance improvement with changing statistic, which was most prominent for the long sound durations (Supplementary Fig. 3). For short duration sounds, there was only a modest improvement in the classifier recognition performance, indicating that the classifier did not effectively make use of the added statistics for very short duration sounds. Regardless of task, it is worthwhile noting that the time constant of the neural spectrum classifier is approximately 200-300 ms faster than that of the neural correlation classifiers. For the +Corr. sound condition, for instance, the time constant is 581 ± 121 ms for the neural correlation and 306 ± 140 ms for the neural spectrum classifier. We also find a similar difference in the discrimination task, where the time constant is 235 ± 109 ms for the neural correlation classifier and 68 ± 14 ms for the neural spectrum classifier for the +Corr. sound condition. However, for the discrimination task, the time constant is relatively stable and does not change substantially with sound statistics (change between conditions Spec. to Ori: 28.8% decrease from 281 ± 101 ms to 200 ± 85 ms for neural correlation; 7.6% from 66 ± 7 ms to 71 ± 22 ms for neural spectrum). Thus overall, texture discrimination can be accomplished by the classifier more quickly than recognition, and spectrum based cues accumulate evidence about the sound more quickly in both tasks.

### Sound Texture Statistics Facilitate Recognition but not Discrimination of Natural Sounds

A series of parallel studies using an identical sound paradigm were carried out to determine how human listeners discriminate and identify sound textures and to determine how different statistics contribute to both tasks. As for the neural classifier, distinct differences were observed in both tasks and distinct performance trends are observed across sound durations and added statistics (Fig. 7A; *p* < 0.001, m-way ANOVA). First, in the recognition task, performance tends to improve upon adding statistics to the synthetic variants (48% improvement from 50.0 ± 10.6% to 98.0 ± 2.7%, Spec. to + Corr. for the 1 s duration; *p* < 0.05, paired t-test) and a significant increase in recognition performance is also observed across sound durations (29% improvement from 71.0 ± 12.9% to 100.0 ± 0.0%, shortest to longest duration for the Ori. condition; *p* < 0.05, paired t-test). Thus, the including high-order statistics seem to have a pervasive role in the subject’s ability to identify the textures used and evidence of these high-order statistics can accumulate over relatively long durations, consistent with prior studies reporting increased perceptual realism with added statistics and sound duration ^1,16^. This behavior sharply contrasts human discrimination trends, where the performance is nearly maxed out and is much more homogenous. Performance improves only subtly upon adding statistics for the short duration sounds (93.0 ± 2.7% to 96.5 ± 3.4%, Spec. to + Corr. for the 62.5 ms duration; *p* < 0.05, paired t-test) whereas it is nearly 100% for all statistics for the longest duration (98.5 ± 1.4% to 99.5 ± 1.0%, Spec. to + Corr. for the 1 s duration; N.S., paired t-test). Thus overall, texture discrimination can be performed relatively quickly requiring few statistical cues and the spectrum cue on its own accounts for most of the discrimination performance. Texture recognition, by comparison, is greatly impacted by adding high-order statistics to the synthetic variants (beyond Spec) and such information can accumulate over time so that recognition performance is highest for the longest sounds.

**Fig. 7:**
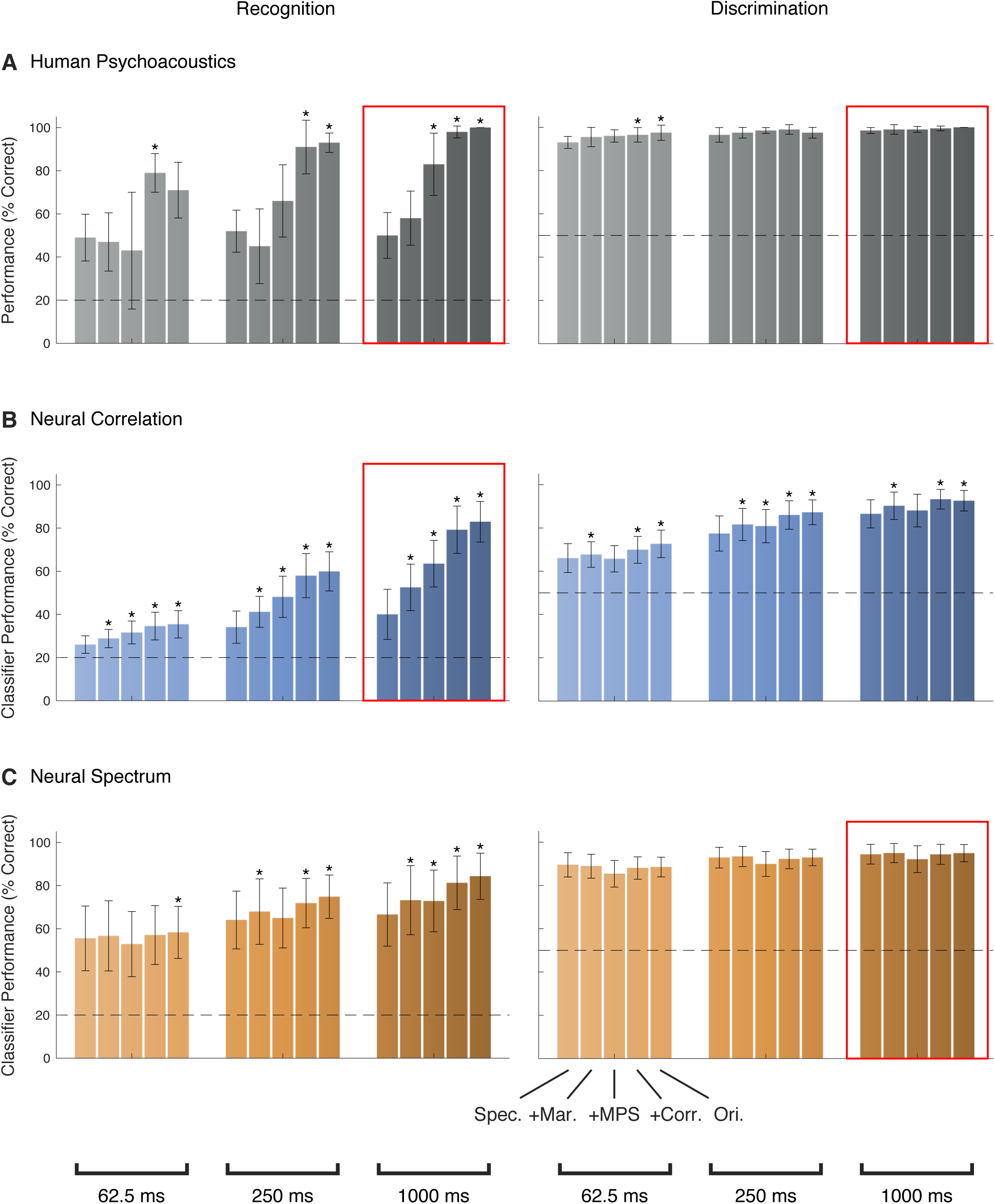
Comparing the task performance between the neural classifiers and human psychoacoustics. (A) Human test scores in the sound recognition (recognition) and discrimination tasks (see METHODS). Results are shown for 62.5, 250, and 1000 ms sound durations and for different summary statistic conditions (Spec., +Mar., +MPS, +Corr., Ori.). (B) Spectro-temporal neural classifier performance as a function of sound statistics in the recognition and discrimination tasks for the same conditions. (C) Neural spectrum classifier performance. Asterisks indicate a statistical significance increase when compared to the Spec. condition (*p* < 0.05, paired t-test, corrected for multiple comparisons).

The human perceptual trends resemble results from the neural classifier where discrimination is fast and recognition slow and where multiple high-order statistics contribute most profoundly to recognition. For this reason, we compared neural classifier against the human listener trends using matched conditions. Fig. 7B and C show the corresponding neural classification trends using spectrum and correlation based neural classifiers. Intriguingly, several parallels between the neural classifier and human trends is observed. First, in the recognition task, performance varied significantly as a function of both duration and added statistics (*p* < 0.001, m-way ANOVA). The neural correlation classifier performance increases substantially with added statistics (39% improvement from 40.0 ± 11.6% to 79.3 ± 11.0%, Spec. to + Corr. for the 1 s duration; *p* < 0.05, paired t-test) and duration (48% improvement from 35.4 ± 6.3% to 83.0 ± 9.4%, shortest to longest duration for the Ori. condition; *p* < 0.05, paired t-test) and the data for 1 s duration follows a similar trend to the human recognition data. The spectrum-based classifier performance likewise shows some improvements with statistics (from 66.6 ± 14.7% to 81.3 ± 12.4%, Spec. to + Corr. for the 1 s duration; *p* < 0.05, paired t-test) and duration (from 58.3 ± 12.1% to 84.3 ± 10.7%, shortest to longest duration for the Ori. condition; *p* < 0.05, paired t-test), but these are more subtle and the performance trends are less similar than for recognition. In contrast, the general agreement between neural classifier and human performance, swap for the discrimination task. Here, the correlation based classifier shows graded improvements with both added statistics (from 86.5 ± 6.5% to 93.3 ± 4.5%, Spec. to + Corr. for the 1 s duration; *p* < 0.05, paired t-test) and (from 72.6 ± 6.4% to 92.6 ± 4.8%, shortest to longest duration for the Ori. condition; *p* < 0.05, paired t-test) which are not observed for human results. By comparison, the spectrum-based classifier follows a nearly identical and much more similar trend to the human data where discrimination performance is nearly maxed out and independent of the duration (from 88.5 ± 4.6% to 94.9 ± 4.0%, shortest to longest duration for the Ori. condition; *p* < 0.05, paired t-test) and statistics included (from 94.4 ± 4.6% to 94.4 ± 4.6%, Spec. to + Corr. for the 1 s duration; N.S., paired t-test). Thus, overall, human perception and neural decoding follow similar trends where discrimination can be accomplished quickly with only spectrum based cues. By comparison, recognition by both human listeners and the neural decoder is much more dependent on high order-statistics and requires longer evidence accumulation times to achieve high performance.

## DISCUSSION

The findings here demonstrate that the spectrum and high-order summary statistics of natural sounds are reflected in the response statistics of auditory midbrain ensembles. Results from the neural decoders are consistent with and predict perceptual patterns for recognition and discrimination in human listeners suggesting that such neural activity can directly mediate auditory behavior. Together the neural and behavioral results support the idea that the statistical structure in sounds is reflected in the neural spectrum and neural correlation statistics of IC neural ensembles and that such neural response statistics serve distinct roles for the recognition and discrimination of natural sounds.

### Neural Construction of Natural Sound Textures

How does the brain construct and represent natural sound textures? The natural textures used in this study are constructed from summary statistics from a relatively simple generative model of the auditory pathway, consisting of a stage of peripheral frequency tuned filters that is followed by a stage of modulation filters. Given the sequential transformation from frequency decomposition in the cochlea to modulation decomposition along the ascending auditory pathway, it is likely that multiple auditory structures contribute to the extraction of summary statistics. Here we have measured three neural response statistics from auditory midbrain ensembles (neural spectrum, temporal and spectral correlation) and shown that these can convey critical sound related information. The auditory midbrain is uniquely situated for representing natural sound summary statistics, given its central position in the ascending auditory pathway where multiple brainstem targets with distinct selectivities converge and where responses from single neurons have been previously shown to be modulated by various natural sound statistics ^4-6,10^. Furthermore, the inferior colliculus is the first stage in the auditory pathway with a preponderance of modulation tuned neurons that ^17^, as a population, could represent the modulation power spectrum of sounds (Mod statistics) and where neurons are uniquely sensitive to correlations across frequency channels ^4,7^. Here we have further demonstrated that the collective statistics from a neural ensemble, not just single neurons, can enable recognition and discrimination of natural sounds textures.

Although the mechanisms underlying the transformation from high-order sound statistics to neural ensemble statistics are not yet clear, specific response statistics may correspond to distinct statistics within the underlying texture synthesis model. For example, the temporal correlations of the neural ensemble are theoretically related to the modulation power spectrum summary statistic (via Fourier Transform), while the spectral correlation reflects synchronous activity across frequency channels, analogous to the Corr summary statistic. There are clear differences between the auditory pathway and the simplified texture synthesis model that preclude a one-to-one mapping between sound and neural response statistics. However, the exact form of the measured statistics may not be critical. More biologically realistic synthesis models with different summary statistics could certainly be selected and might more accurately predict behavior or the neural activity. More important to our conclusions, is the general observation that statistics of the sound modulate neural response statistics and that statistics beyond the spectra appear to influence neural coding and perception.

### Relationship to Visual Textures

Texture synthesis methods were originally developed for vision ^18^, and both visual and auditory texture synthesis models involve similar receptive field hierarchies and similar statistics (power spectrum, marginals, correlations) ^1,18^. As for audition, natural scene statistics from such models can drive the firing of visual cortex neurons in ways that likely contribute to visual perception ^19,20^. Furthermore, similar to what we report here for IC, correlated firing in visual cortex can be driven directly by high-order visual scene structure ^21^.

The recent use of texture synthesis to study vision and audition ^16,18^ is in part motivated by the fact that realistic texture stimuli can be generated with models that account for underlying transformations of both modalities and which suggest common sensory processing principles. However, there are some noticeable difference between the two modalities. Perhaps the most noticeable distinction is the fact that auditory textures overwhelmingly rely on temporal sound structure whereas visual textures do not. Visual textures can be generated with a purely static model (no time dependency) which contains spatially segregated receptive fields and summary statistics that account for the high-order spatial statistics between image pixels. By comparison, natural sound texture synthesis involves receptive fields that operate simultaneously in time and frequency and where all of the measured summary statistics are averaged over time. While most of the relevant information in dynamic natural scenes is relatively slow (< 10 Hz) ^22^, natural sounds contain perceptually salient temporal cues which cover several orders of magnitude (∼0.5 Hz −1.0 KHz) ^23^. These cues span distinct perceptual ranges of rhythm, roughness, and pitch and auditory neurons selectively synchronize to these ranges across the auditory hierarchy ^23,24^. Thus, in contrast to vision, where textures can be identified with purely statistic images that contain spatial statistics only ^25^, natural sound textures require temporal structure to be meaningful ^1^.

As shown here, many of the informative high-order neural response statistics are inherently temporal. For instance, the spectral and temporal correlation statistics are both derived from temporal response that are synchronized to the sound envelopes up to several hundredths of Hz and thus both statistics rely heavily on temporal synchronized neural activity. Furthermore, a static sound (constant spectrum, as for Spec) that lacks temporal structure, does not form a realistic auditory impression ^1^ and, as shown here, cannot be easily identified as arising from a unique auditory source by humans or the neural decoders. Thus, a major advantage of texture stimuli is that they have modifiable high-order structure which is critical perceptually. As shown, neural responses are sensitive to this structure and, given the highly nonlinear transformations in both vision and audition, these stimuli may lead to a more complete understanding of perception compared to simpler frequency-based stimuli, such as tones or amplitude modulated sounds.

### Comparing Perception and Neural Decoding

Neural response statistics in IC ensembles reflect the summary sound statistics which enabled both discrimination and recognition of the natural sound textures studied here. Adding summary statistics to the synthetic texture variants produces neural correlations that more closely resemble that of the original sound and which are more easily identified. Similar behavior was observed for both the human listeners and neural data whereby the classification performance improves upon adding summary statistics to the synthetic variants. Although classifier performance improved when using either neural spectrum or correlation statistics, the neural spectrum does not vary substantially when adding summary statistics, despite large perceptual differences in these sounds. Moreover, the recognition performance improvements were larger for correlation statistics and the trends more closely resemble human perceptual trends (Fig. 7). This is partly expected, since neural correlation statistics predict and are themselves correlated with the summary statistics of the sound ^4^.

While texture recognition appears to be linked to high-order sound and response structure, texture discrimination is much more stable and appears to depend much less on which summary statistics are included. One notable difference between neural classification and human psychoacoustical data is that the neural classifier is trained using the five original textures sounds for the recognition task, whereas humans were presented these sounds blindly without feedback. Thus, human listeners have no knowledge of the particular statistics for the five textures sounds and instead they relied on prior learned knowledge for these sound categories. Despite this, both humans and the neural classifier appear to be able to utilize summary statistics for recognition since, in both instances, adding statistics to the synthetic variants improves performance. In contrast, the neural discrimination classifier used the neural response statistics directly from the two sounds being discriminated which more closely resembles the perceptual comparison between the two sounds carried out by human listeners. Sounds could be easily discriminated, both by human observers and the neural classifier, even if high-order summary statistics are not included in the synthetic variants. This suggests that the sound spectrum, while not the sole cue used for discrimination, may be sufficient for discrimination on its own. Spectrum equalized sounds can be discriminated and recognized from neural ensemble responses (Fig. 5D). Moreover, these sounds are perceptually quite different and can be readily identified or distinguished from each other (Supplementary sounds 26-30). Thus, if spectrum cues are not available it is possible to take advantage of high-order structure in the neural activity (neural correlations) for both recognition and discrimination.

An intriguing aspect of both the neural and human findings is that the evidence integration time scales can depend critically on both the chosen statistic and the perceptual task. Texture discrimination can be accomplished relatively easily with spectrum based cues only and, with both the human listeners and neural decoders, and the integration of spectrum cues is exceptionally fast. Although this does not exclude the possibility of high-order cues being used, it suggests that spectrum cues are sufficient for high accuracy natural texture discrimination. This is consistent with prior findings suggesting that texture excerpts (different exemplars of the same texture) can be discriminated readily with short duration sounds where spectrum cues dominate and that they become increasingly difficult to discriminate for longer durations, where presumably the average spectrum and high-order statistics are similar ^16^. By comparison, our data suggests that recognition of natural sound textures appears to depend much more heavily on the availability of high-order summary statistics and evidence about the summary statistics need to be accumulated over relatively long durations to be informative, both neurally and behaviorally. Such findings are consistent with prior perceptual studies where it has been shown that high-order summary statistics are necessary for creating realistic impressions of sounds ^1^ and that texture discrimination (from different categories) improves with increasing sound duration ^16^. Our results build on these findings by suggesting that low- and high-order summary statistics appear to have distinct evidence integration times and these appear to contribute differently to recognition and discrimination of sound textures.

Altogether, the findings here suggest that information from neural response statistics contribute differentially to recognition and discrimination of sound textures. Low-order summary statistics (e.g., spectrum) and the corresponding neural response statistics (e.g., neural spectrum), accumulate information quickly allowing for fast and accurate sound discrimination. High-order sound statistics, by comparison, are reflected in coordinated activity across IC neural ensembles (neural correlations). Such activity requires longer evidence accumulation times to be useful yet it contributes substantially more towards the recognition of natural sounds.

## METHODS

### Sound Statistics and Synthesis

Five natural sound textures were used in this study: crackling fire, bird chorus, outdoor crowd, running water, and rattling snake sounds. These sounds were selected since they each have distinct spectral and temporal properties. These real-world textures were initially analyzed in a generative auditory model that contained hierarchical filters representing the signal processing of the cochlea and mid-level (e.g. auditory midbrain) auditory system, and the statistics of the resulting decomposition were measured ^1^.

Briefly, in the cochlear model stage the sound waveform was processed through 32 tonotopically arranged cochlear filters (100 Hz – 20 kHz), followed by nonlinear compression and lowpass filtering to extract the envelopes of each cochlear model channel. We refer to the resulting output as a cochleogram (cochlea spectrogram, Fig. 1A) since it depicts the time and frequency envelope fluctuations at the output of the cochlear model. The mid-level auditory model consisted of an ordered modulation filterbank (20 logarithmically spaced filters between 0.5-200 Hz) which models the modulation decomposition of sounds carried out by the central auditory system.

Next, for each texture sound we measured several statistics directly from the cochlear and mid-level outputs of the model. These statistics were used to characterize the sound structure and to generate synthetic variants with perturbed statistics. These perturbed variants were then use to study how physiologically guided statistics are encoded and how they contribute to sound recognition and discrimination. The cochlear model output statistics included the cochlear channel power and cochlear envelope marginals (mean, variance and skewness of the cochleogram envelopes., Fig. 1B) and the correlation matrix between distinct frequency channels Fig. 1D), which we refer to as the spectral correlations. At the mid-level stage, we computed the modulation power spectrum (Fig. 1C), which depicts the power distribution of the envelope fluctuations as a function of modulation frequency. In addition, we also computed the second-order correlations between modulation channels.

The measured texture statistics from the auditory model were then used for sound synthesis. These perturbed sounds were used for the animal physiology and human psychoacoustics described below. Synthetic sound variants were initialized with a Gaussian white noise signal, an iterative procedure was used to modify the signal until it included the desired statistics ^1^. In our study, for each original sound texture, we first generated corresponding synthetic sound variants with only the cochlea spectrum (cochlea channel power) imposed. Additional variants were then created which we sequentially incorporated the cochlear marginal, modulation power spectrum, and correlations. For simplicity, we refer to these synthetic variants as Spec., +Mar., +MPS and +Corr. conditions throughout the manuscript. Perceptually, the Spec. only sounds were perceived as frequency-shaped noise, and adding additional statistics created variants that were more realistic. The complete model variants were nearly indistinguishable from the original sounds.

In a control study, we generated sound variants with a matched 1/f power spectrum (pink noise) ^4^. This manipulation removed all spectral cues while preserving many of the fine structure and high-order modulation cues in the original sounds. To synthesize such spectrum-equalized sounds, the sound magnitude spectrum S(*f*) (obtained using a Welch average periodogram) was used to generate a zero-phase inversion filter with a transfer function of

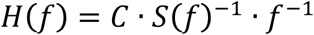

where *C* is a gain normalization constant to control the sound pressure levels (SPLs). A synthetic sound variant with a 1/f spectrum was then generated by filtering the sound with *H* (*f*).

### Animal Sound Delivery

Sound stimuli were delivered to the animal via custom fitted ear molds (see Animal Procedures and Surgery) that contained a sound delivery tube and attached to a closed binaural speaker system (Beyer DT 770). The sound delivery system was calibrated (flat spectrum between 0.1 and 24 kHz, ± 3 dB) for each animal with a finite impulse response inverse filter, and implemented with a Tucker-Davis Technologies (TDT) RX6 multifunction processor.

To evaluate whether the recording location was within the central nucleus of the inferior colliculus (IC), we first presented a random sequence of pure tones (98 kHz sampling rate, 50 ms duration tone pips with 5 ms cosine-squared ramp and 300 ms inter-tone interval, spanning 0.1-20 kHz and 5-75 dB sound pressure level (SPL)) to measure the frequency response area of each recording channel. This allowed us to estimate the frequency selectivity of each penetration site, and we only include locations that showed a tonotopic gradient in further analysis.

Next, we delivered a sound texture paradigm which consisted of the original textures sounds along with perturbed synthetic variants. The five original sound textures, including fire, bird, crowd, water and snake, along with their synthetic variants generated using texture synthesis by sequentially including the spectrum, marginal, modulation power and correlations statistics were delivered in a block randomized fashion (3 s sound duration, 100 ms inter-stimulus interval). The sound stimuli were delivered at 65 dB SPL for 20 to 50 mins (17 to 39 trials) depending on the recording stability.

In the spectrum-equalized paradigm, the five 1/f equalized sounds were interleaved with the five original sounds and delivered in a similar block randomized fashion as described above at 65 dB SPL. Sounds were delivered for 20 trials in all recording sites except for one site, which had 14 trials.

### Animal Procedures and Surgery

In this study, four female Dutch-Belted rabbits (age of 0.5-2 years) with a weight of 1.5-2.5 kg were used. We measured the auditory response properties of neuron ensembles in the auditory midbrain (inferior colliculus, IC) of unanesthetized animals during passive listening to experimental sounds. All experimental procedures were approved by the University of Connecticut Animal Care and Use Committee and in accordance with National Institutes of Health and the American Veterinary Medical Association guidelines.

Animals were initially trained to sit still for the desired duration of an experiment session. Then surgical procedures were carried out in two phases with a recovery and acclimation period between procedures. All surgical procedures were performed using aseptic techniques. Animals were first sedated with acepromazine, and deep levels of anesthesia were maintained with isoflurane (1-3%) and oxygen (1 liters/min) throughout the surgery, while electrocardiogram and rectal temperature were monitored. The first procedure involved a surgical implantation of a head restraint bar, which assured stable neuronal recordings without head movement, as well as the fabrication of custom ear molds, which were used for sound delivery. First, the procedure involves a midline incision on the scalp to expose the sagittal suture between bregma and 2-3 mm posterior to lambda. Stainless steel screws (0-80) and dental acrylic were used to affix a brass restraint bar oriented rostro-caudally and to the left of the sagittal suture. Dental acrylic was then used to form a dam on the exposed skull on the right hemisphere between bregma and lambda. Next, a small cotton ball was inserted to block the external auditory meatus, and a vinyl polysiloxane impression material (VPS) was poured into the animal’s ear canals. After hardening, the impression compound was removed and it was subsequently used to build a cast from which custom ear molds were fabricated for each ear.

Following the first surgery and a 5-day postoperative recovery period, an acclimation was conducted for about two weeks during which the animal was trained to sit still with the head restraint while sounds were delivered via the ear molds. Once the animal was habituated to the sound delivery, the second surgical procedure, which consisted of a small craniotomy, was performed. An opening (approximately 4 × 4 mm) was made on the right hemisphere within the dental acrylic dam, and centered approximately 12-13 mm posterior to bregma. The brain area was then sterilized with disinfectant and antiseptic (chlorhexidine solution), and vinyl polysiloxane material was applied into the acrylic dam to cover and seal the exposed region.

### Multichannel Neurophysiological Recordings

Animal training and experiment recordings were conducted in a double-wall sound-attenuation acoustic booth (Industrial Acoustic Company). Multi-channel silicon probes (NeuroNexus, 16-channel linear electrode arrays, 32-channel polytrodes, and 64-channel polytrodes with electrode separation of 150, 50 and 60 um, respectively, 1-3 MΩ impedance at 1 kHz) were used for acute extracellular neurophysiological recordings in the inferior colliculus (IC) of unanesthetized rabbits. We selected a subset of 16 equally spaced channels from the 32 and 64-channel polytrode recordings to maintain a uniform IC sampling and data format for analysis. During each recording session, the electrode probe was advanced through the dura into the brain using a micromanipulator (Burleigh EXFO LS6000) to a depth of approximately 7.5-9.5 mm relative to the cortical surface at which clear responses to brief bursts of broadband noise or tones could be seen from the majority of the electrodes. During sound delivery, neural activity was acquired continuously at a sampling rate of 12 kHz a PZ2 preamplifier and a RZ2 real time processor (TDT).

For the sound texture paradigm, we recorded from a total of 38 penetration sites in 4 animals and 29 sites were used in this study due to the data quality based on response stability, recording duration (a minimum of 15 trials per sound variant), and the covered frequency range (at least two octaves). For the spectrum equalized paradigm, we recorded from an additional 11 penetration sites from 2 animals.

The recorded neural traces were analyzed offline using custom MATLAB (MathWorks) codes. We used an analog representation of multi-unit activity (aMUA) for analysis as opposed to single unit or thresholded MUA (tMUA) activity for several reasons. First, to study the neural population activity across the frequency span of IC we needed to densely sample the tonotopic axis of IC, which is not possible with single units. Using spike sorting, we can typically identify ∼2 to 5 single units per recording session with the 64 channel probes, which is a relatively sparse sampling of the neural activity along the frequency axis. Although we could use tMUA, previous studies have shown that aMUA captures the structure of population activity with substantially less noise, and it provided similar neural responses when compared to tMUA ^4,26^. To obtain aMUA, the recorded voltage signals of each electrode channel were first bandpass filtered to 325-3000 Hz (b-spline filter 125 Hz transition width, 60 dB attenuation), and then full-wave rectified and lowpass filtered with a cutoff frequency of 475 Hz (b-spline filter 125 Hz transition width, 60 dB attenuation). The resulting envelop signals were further down-sampled to 2 kHz for computation efficiency. A data raster was generated for each electrode channel using such aMUA envelope signals over time and across recording trials (Fig 2B).

### Population Response Metrics: Neural Correlations and Neural Spectrum

We separately assessed the role of spectrum- and correlation-based codes and their contribution towards neural recognition and discrimination of natural sound textures. Spectrum-based codes, account for the tonotopic decomposition of sounds along the auditory pathway. It can be viewed as a conventional place-rate code where the strength of activity is driven bottom up by the power in the sound at each frequency channel. Here, the neural spectrum for a given sound at a specific recording location is estimated by averaging the responses from each channel across trials and time

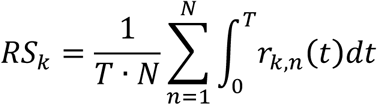

where *r*_*k,n*_ (*t*) is the *n*^th^ response trial from channel *l*. Conceptually, the neural spectrum accounts for the frequency specific strength of firing across the recording array channels (Fig. 2E).

Neurons in IC can potentially also encode stimulus information through coordinated firing across the neural population, and for this reason we also investigated a correlation-based codes in which pairs of neurons at different spatial locations can show synchronous activity. We estimated the **neural correlations** (both *stimulus-driven* and *noise correlations*) across the 16 recording channels. The analysis procedure used in this study is identical to that described in a recent publication ^4^, where the cross-channel correlations were “shuffled” ^15,27^ across response trials and they were computed with a short time window ^28^. Eliminating the trial-to-trial variability of the neural activity, this windowed shuffled correlation allowed us to isolate stimulus-driven correlations from the responses. For a pair of spatially separated electrode channels *k* and *l*, the short-term correlation was computed as:

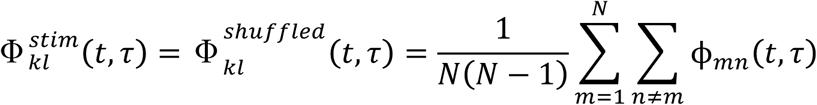

where ϕ_*kl,mn*_(*t*, τ) is the windowed cross-correlation between the *m*^th^ and *n*^th^ response trials, and *N*(*N-*1) is the total number of trial pairs that are correlated. The correlation at time *t* with delay *τ* was given as:

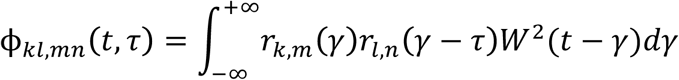

where. *r*_*k,m*_ (*t*) is the signal (mean removed) in the *m*^th^ response trial from channel *k* and *r*_*l,n*_(*t*) is the signal (mean removed) in the *n*^th^ response trial from channel *l*, while *γ* is a time integration variable. *w*(*γ*) is a unit amplitude centering square window applied about *γ*=0 with a duration of T (varied between 62.5 to 1000 ms). The above shuffled correlations were implemented using a fast algorithm as outlined previously ^4,27^.

The short-term correlation was further normalized as a correlation coefficient. This version allows us to measure stimulus driven correlated activity irrespective of the response power for each recording channel. For this case, the signal variance depends on the selected time and delay samples used for the above correlation estimates. Thus, the short-term signal variance was estimated across time and delay samples and the expectation was computed by averaging over trials as:

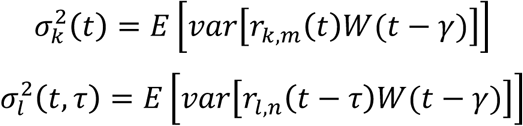

Then the normalized correlation is then obtained as:

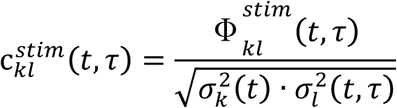

where 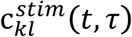; is the correlation coefficient between -1 and 1.

To assess the contribution of nonspecific neural activity, we also estimated noise correlations, which are the result of stimulus-independent network activity. The noise correlations were obtained by subtracting the shuffled (different trials) from the unshuffled (same trials) cross-correlations ^4^. Similar to the shuffled correlation, the unshuffled correlation between recording channels *k* and *l* was given as:

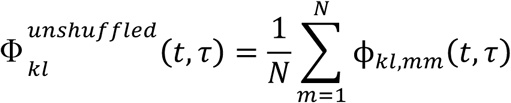

where we note that identical trials (*m*) is used to correlate the activity of different channels (*k* vs. *l*). The unnormalized and normalized noise correlations are then obtained as:

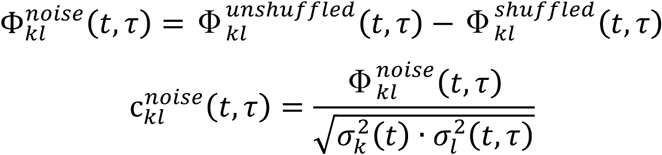

where 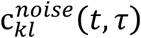 is also bounded between −1 and 1. Collectively, the total neural correlation is obtained by combining the stimulus-driven and noise correlations at zero lag were 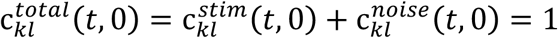.

As described above, the correlations were measured between spatially separated electrode channels (e.g. channels *k* and *l*) using aMUA envelopes. Due to the fact that IC is tonotopically organized, our recordings of the electrode channels are also frequency ordered (Fig. 2A). Thus, we refer to spatial correlations as spectral correlations. The description of **spectral correlations** in this work specifically means zero-lag correlations between electrode channels, c_*spec*_ (*t*) = c_*kl*_ (*t*, 0) (Fig. 2C). By considering the correlations between the same recording channels (e.g. the diagonal of Fig. 2C, channel *k*=*l*) at different delays, we also obtained a measure of **temporal correlations**, c_*temp*_ (*t*) *h* c_*kk*_ (*t*, τ) with *τ* extending from −100 to 100 ms (Fig. 2D). Fig. 2D and F show a segment of aMUA envelopes from 4 electrode channels with their pairwise scatter plots in a representative penetration site.

### Assessing Neural Population Activity as a Function of Sound Statistics

We also quantified how the neural population responses change upon adding stimulus statistics to the synthetic sound variants and asked how incorporating higher-order structure in the texture sounds alters neural responses. We considered each of three-response metrics (neural spectrum, spectral correlation, or temporal correlation) separately and quantified how these differed from the original sound responses. First, we treated each of the response metrics as a vector for the synthetic and original sound texture responses

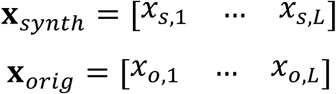

where *X*_*synth*_ is the vectorized response metric for the synthetic condition obtained for the first half of the data and *X*_*orig*_ is the vectorized response for the original sound texture obtained from the second half of the data. To asses the similarity of the response pattern, we computed a similarity index between the synthetic and original responses as

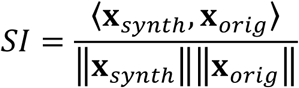

where it is equivalent to a correlation coefficient between the synthetic and original response and −1 ≤ *SI* ≤ 1. Note that in the instance where the synthetic condition used for comparison is the original sound response, the above SI quantifies the similarity of the 1^st^ and 2^nd^ response halves for the original sound, thus setting an upper bound on the SI, which is limited by measurement noise.

Next, we quantified how the strength of each of the response metrics (spectrum or correlation) varies with changing statistics of the synthetic variants. To do so, we defined that strength ratio between the synthetic and original sound response as

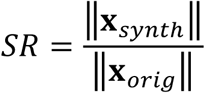

where ‖ · ‖ represents the Euclidean norm operator. The *SR* assumes values 0 < *SR* < 1 whenever the synthetic response is weaker than the corresponding original sound response. Values of *SR* > 1 are obtained if the synthetic response is stronger than the corresponding original sound response. In general, we found that the synthetic sound responses where typically weaker than those of the original sound so that the measured *SR* values where always between 0 and 1.

### Single-Trial Neural Classifiers

To evaluate the extent to which the neural spectrum and correlation statistics contribute to recognition and discrimination of texture sounds, we developed a single trial neural classifier which we applied separately in texture recognition and texture discrimination paradigms. The classifier has been described in detail previously ^4^. Briefly, we use a cross-validated naïve Bayes classifier using a low dimensional representation of the neural correlations or the neural spectrum as features. We first applied principal component (PC) analysis to the measured response feature vectors to obtain a low dimensional representation of the IC neural activity. A multiclass naïve Bayes model was then trained using the low dimensional PC representation where the class labels correspond to individual textures (crackling fire, bird chorus, outdoor crowd, running water, and rattling snake). Each of the PC score distributions was fit to an axis-aligned multivariate Gaussian distribution and the number of PCs used for the multivariate distributions was chosen to maximize the cross validated performance across all recordings (number of PCs chosen vary from 4 to 18 depends on the type of classifier and task analyzed).

Next, given an input single trial neural response of specified duration the maximum a posteriori (MAP) rule was used to classify and assign a class label (for details, see ^4^). To test for overfitting, the model was optimized using the first half of neural response data for training and the second half for validation. This procedure was then repeated by swapping the first and second half of the data. Since we are interested in investigating the classification performance as a function of sound duration (or equivalently, neural response duration), different data segments were selected from the training and/or validation data set with a window length that was varied from 62.5 ms to 1000 ms in ½ octave steps. At each window length, we selected response measurements centered about 100 randomly selected locations.

To identify the roles that different neural *response statistics* might play in discrimination or recognition, the classifier was applied separately to the neural spectrum, temporal correlation, spectral correlation and the joint spectro-temporal correlations as described previously ^4^. Furthermore, we were interested in characterizing how the *sound statistics* potentially influence recognition and discrimination, and, for this reason, the model was validated using responses from both the original sounds and all of the synthetic variants with reduced statistics.

For each of the neural response metrics and synthetic sound variants, the neural classifier was applied separately for the texture recognition and discrimination paradigms. In the case of texture recognition, the classifier was required to identify a sound from the provided neural response in a five-alternative forced choice task. In this case, the goal is to identify the neural activity that most closely resembles that of the original sound, and thus the model for each of the original five texture sounds was used (using 1000 ms window). The classification rule was evaluated for each of the five sound models given the provided neural response (either from synthetic or original variants) and the class assignment was based on maximizing the posterior probability. For the sound discrimination paradigm, the goal was to determine whether the neural responses to two provided sounds (at a specified sound duration) could be differentiated from one another. In this case, in addition to having identical duration, both sounds tested were selected from variants for the same statistic condition (synthetic or original; e.g., fire vs. bird sounds both containing +MPS condition). For the purpose of discriminating the sounds using neural activity, the models derived were generated using the neural responses for the same two sounds using the matched statistic condition (e.g. fire and bird sounds with +MPS condition). We then use classifier to estimate which of the two sounds was delivered using the MAP rule. Note that because we are now comparing against two possible sound textures, the performance will be at chance (50%) whenever the two sounds are not discriminable. The classifiers were implemented in a similar fashion for the spectrum-equalized paradigm. In the case of recognition, the model was trained using the responses to the original sounds, and the validation data were obtained from the responses to the 1/f spectrum sounds. In the case of discrimination, both training and testing were carried out using the data of 1/f spectrum sounds.

### Human Psychoacoustics

We carried out complementary experiments in human subjects to determine how different sound statistics contribute to recognition and discrimination of texture sounds. All procedures for these studies were approved by the Institutional Review Board at the University of Connecticut.

The original sounds used for these studies are the same as those used in the animal physiology. We recruited 2 male and 3 female subjects ages 20-35 with normal hearing for the experiments. Subjects sat in a single-walled sound attenuation chamber (Industrial Acoustic Company) and were asked to identify or discriminate both original and synthetic texture sounds by responding on a keyboard. Each paradigm was broken up in blocks of approximately 5 min and subjects were allowed a rest period of 15 min midway through the experiment session. All sounds were delivered at a 44.1 kHz sampling rate using an RME Fireface UC digital audio interface (RME Audio) through ER-4 in-ear insert headphones (Etymotic Research).

In the sound recognition paradigm, subjects performed a five-alternative forced choice task in which they were required to determine the identity of the sound delivered. The sounds used and the corresponding response labels were the same as the sounds used in the animal physiology. Subjects reported the sound heard by pressing one of five labeled keys on a keyboard (fire, water, birds, crowd, snake) with a pictorial icon for each sound situated above each key. Since the goal is to determine how different statistics contribute to recognition and the integration time-scales at which recognition takes place, synthetic and original sound variants were delivered for three different sound durations (62.5, 250, and 1000 ms) and all sounds were delivered for 4 trials. To minimize the possibility that subjects learn or memorize a particular sound features unique to any of the textures and instead focus on the overall summary statistics, sounds were delivered in a hierarchical blocked order were within each main block, the sounds testing progressed sequentially in sub-blocks from shortest to the longest sound durations. For a given duration sub-block, we delivered all five sounds and trials in randomized order. Sequential test blocks, on the other hand, were used to add statistics. That is, we first tested all subjects on the Spec. condition, followed sequentially by +Mar., then +MPS, +Corr., and Ori. Within a block, only one statistic was tested and all of the 62.5 ms sounds were tested first in random order, followed by 250 ms and 1000 ms. These controls were necessary to minimize the possibility that subjects learn and memorize the unique sound spectrum and possible high-order features for each sound, particularly for short duration sounds.

In the discrimination paradigm, subjects performed a two-alternative forced choice tasks in which two sounds are delivered and subjects are required to respond whether the two sounds are the same or different. As for the recognition paradigm, the sounds and statistics used, the block ordering for different statistics, and sound durations are identical. However, whereas in the recognition paradigm only one sound is delivered, every trial now has two randomly selected sounds for each trial. All trials are matched so that the statistics used and durations of the delivered sounds are identical. However, half of the trials contained randomly selected variants of the same texture (fire vs. fire) while the remaining half contained randomly selected variants from the different texture sounds (e.g., fire vs. water). Thus, chance performance for all of the tested conditions is at 50%.

## Acknowledgements

We thank the late Dr. S. Kuwada for generous support and guidance on experimental procedures in the unanesthetized rabbit.

---

**Supplementary Figure 1:**
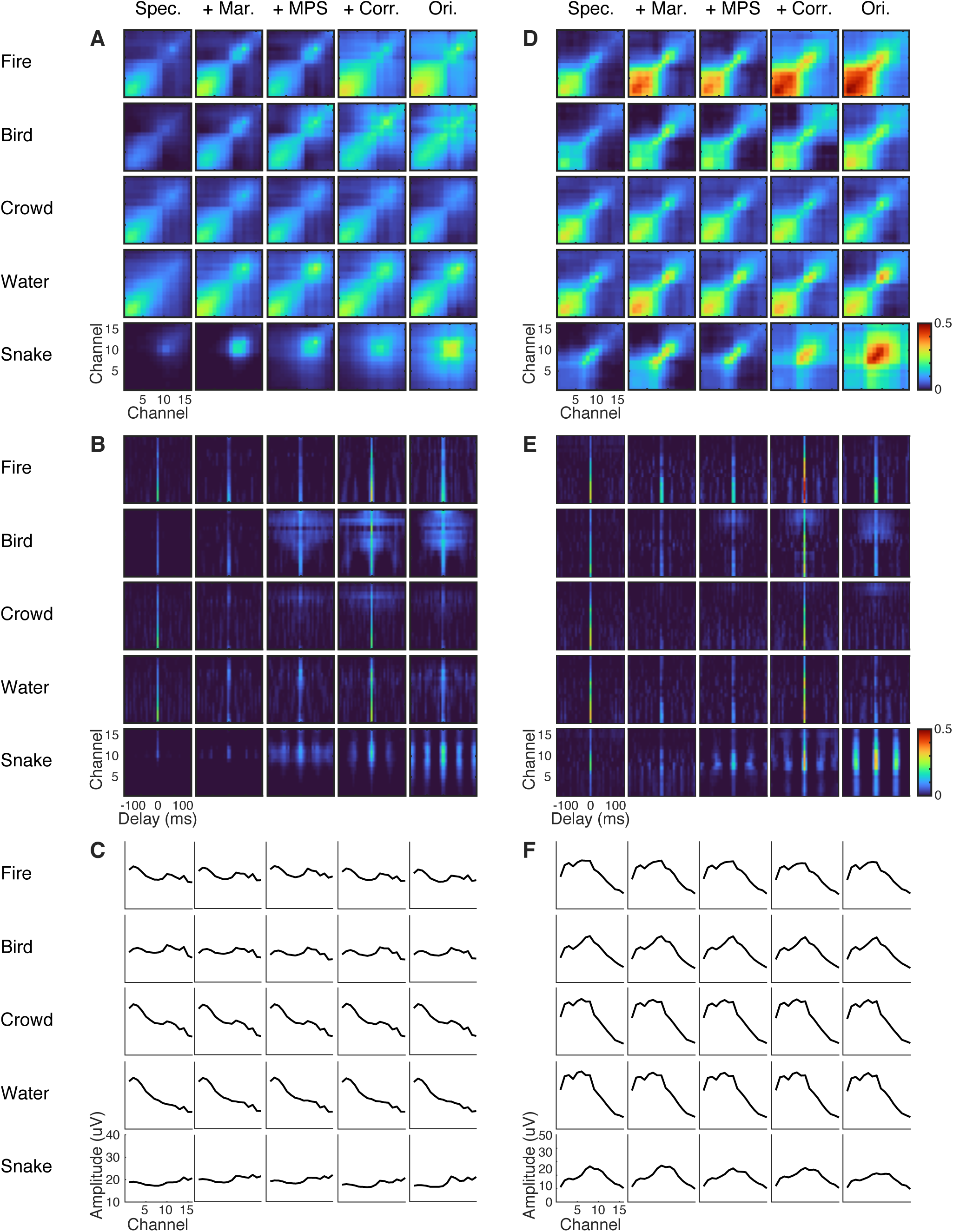
Neural correlations and response spectrums for synthetic variants and original sounds in two example penetration sites. The data is formated as for Fig. 3A, B and C.

**Supplementary Figure 2:**
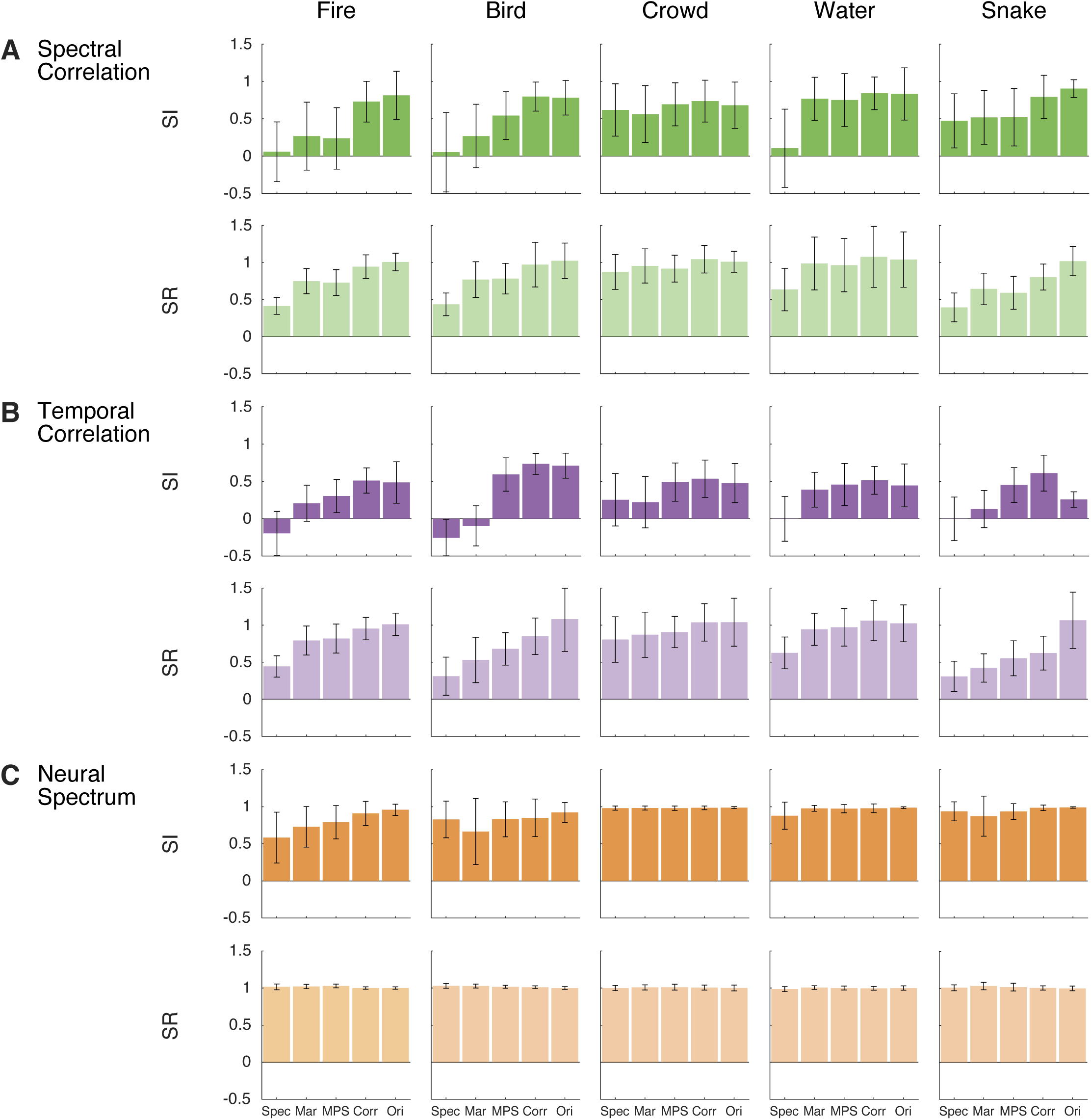
Average similarity index (SI) and strength ratio (SR) as a function of sound statistics across *N*=29 penetration sites (*N*=4, 9, 8, 8 from four animals) for five different sounds. (A-B) For each sound, the averaged similarity index and relative strength of spectral and temporal correlations increase with statistics, indicating the correlation matrices converge upon adding sound statistics. (C) Using response spectrum, both similarity index and relative strength do not vary substantially with sound statistics in all conditions except the similarity index of the fire sound increases from 0.58 to 0.96. Error bars, S.D.

**Supplementary Figure 3:**
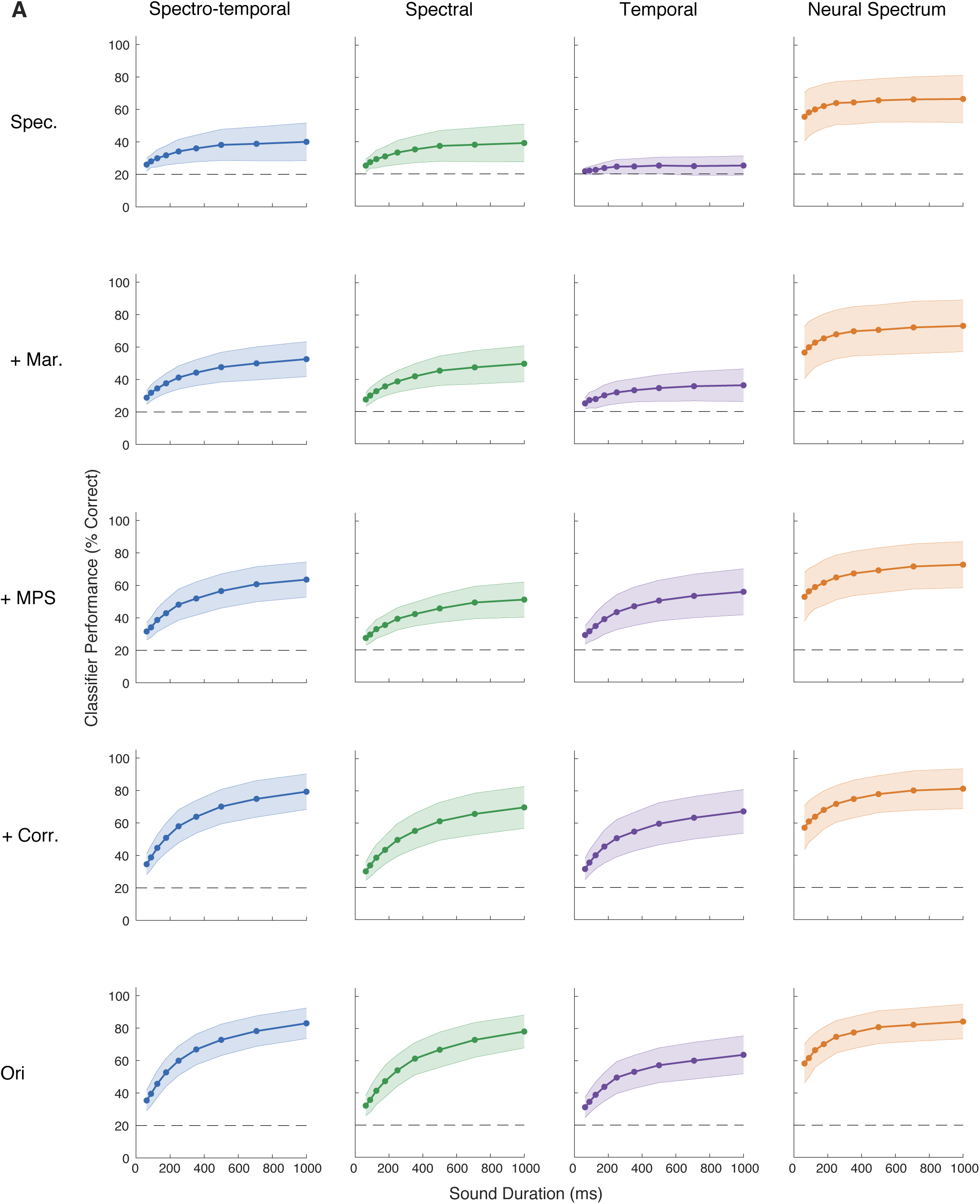
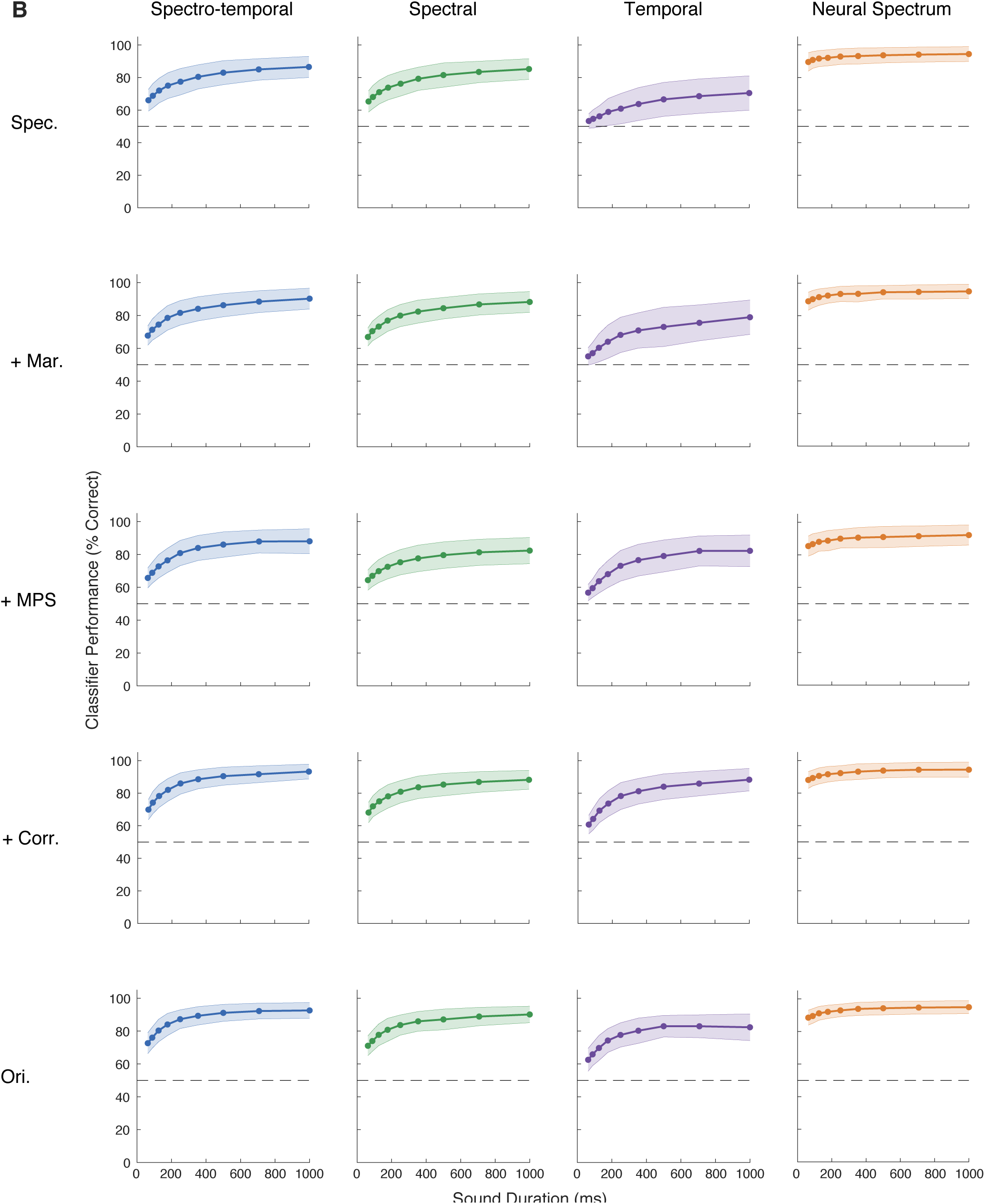
Average classification performance across *N*=29 penetration sites (*N*=4, 9, 8, 8 from four animals) under different sound statistic conditions. (A) Sound identification task. (B) Sound discrimination task. Shaded areas, S.D.

**Supplementary Figure 4:**
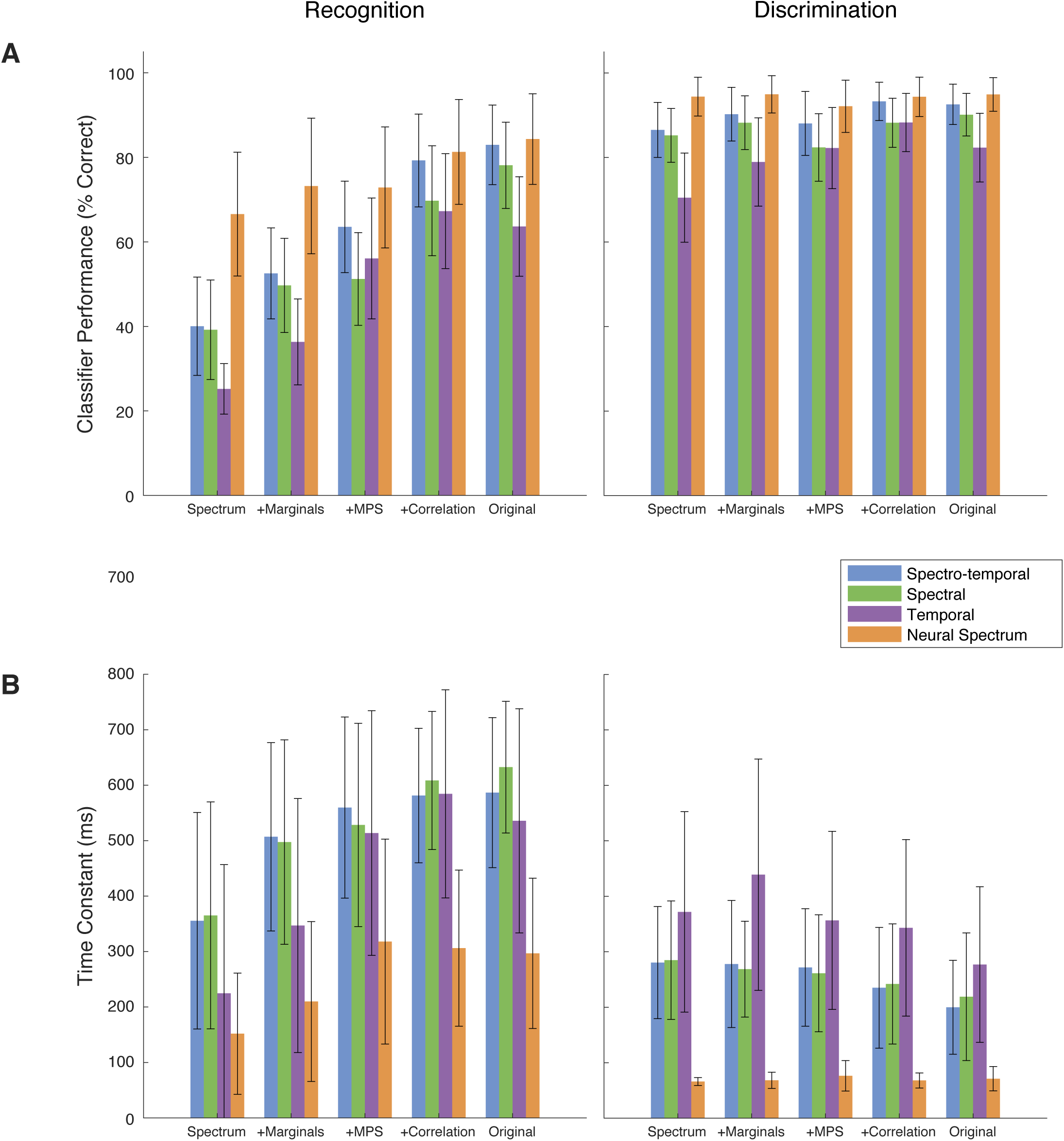
Neural classifier performance and time constant as a function of sound statistics. (A) Classification results for the spectro-temporal, spectral, temporal, and response spectrum classifiers in the identification and discrimination tasks. Shown are the average performances (across sounds and penetration sites) with 1 s sound duration. (B) Time constant (time taken to reach 90% of the corresponding maximum performance) as a function of sound statistics in the identification and discrimination tasks. Error bars, S.D.

